# *Sry*-modified laboratory rat lines to study sex-chromosome effects underlying sex differences in physiology and disease: Four Core Genotypes and more

**DOI:** 10.1101/2023.02.09.527738

**Authors:** Arthur P. Arnold, Xuqi Chen, Michael N. Grzybowski, Janelle M. Ryan, Dale R. Sengelaub, Tara Mohanroy, V. Andree Furlan, Helen R. Schmidtke, Jeremy W. Prokop, Monika Tutaj, William Grisham, Shanie Landen, Lynn Malloy, Akiko Takizawa, Julia L. Ciosek, Kai Li, Theodore S. Kalbfleisch, Hayk Barseghyan, Carrie B. Wiese, Laurent Vergnes, Karen Reue, Jonathan Wanagat, Helen Skaletsky, David C. Page, Vincent R. Harley, Melinda R. Dwinell, Aron M. Geurts

**Affiliations:** Department of Integrative Biology & Physiology, and Laboratory of Neuroendocrinology of the Brain Research Institute, University of California, 450 Hilgard Avenue, Los Angeles, Los Angeles 90095 CA USA; Department of Physiology, Medical College of Wisconsin, Milwaukee, WI 53226, USA; Hudson Institute of Medical Research, Monash Medical Centre, Melbourne, Australia; Department of Psychological and Brain Sciences, Indiana University, Bloomington, IN, USA; Office of Research, Corewell Health, Grand Rapids, MI 49503 USA; Department of Psychology, University of California, 405 Hilgard Avenue, Los Angeles, Los Angeles, CA 90095, USA; Department of Veterinary Science, Martin-Gatton College of Agriculture, Food, and Environment, University of Kentucky, Lexington, Kentucky 40546, USA; Center for Precision Medicine and Genomics Research, Children’s National Hospital, Washington, District of Columbia, USA; Department of Human Genetics, David Geffen School of Medicine at UCLA, Los Angeles, 90095 CA, USA; Department of Medicine, Division of Geriatrics, David Geffen School of Medicine at UCLA, Los Angeles, 90095 CA, USA; Howard Hughes Medical Institute, Whitehead Institute, Cambridge MA, USA

**Keywords:** *Sry*, testis determination, sexual differentiation, sex determination, sex-chromosome effect, Four Core Genotypes, X chromosome, Y chromosome

## Abstract

**Background:** Previous research on Four Core Genotypes and XY* mice has been instrumental in establishing important effects of sex-chromosome complement that cause sex differences in physiology and disease. We have generated rat models using similar modifications of the testis-determining gene *Sry*, to produce XX and XY rats with the same type of gonad, as well as XO, XXY and XYY rats with varying gonads. The models permit discovery of novel sex-chromosome effects (XX vs. XY) that contribute to sex differences in any rat phenotype, and test for effects of different numbers of X or Y chromosomes.

**Methods:** XY rats were created with an autosomal transgene of *Sry*, producing XX and XY progeny with testes. In other rats, CRISPR-Cas9 technology was used to remove Y chromosome factors that initiate testis differentiation, producing fertile XY gonadal females. Interbreeding of these lines produced rats with interesting combinations of sex chromosomes and gonads: XO, XX, XY, XXY rats with ovaries; and XO, XX, XY, XXY, and XYY rats with testes. These groups can be compared to detect sex differences caused by sex-chromosome complement (XX vs. XY) and/or by gonadal hormones (rats with testes vs. ovaries). Other comparisons detect the effects of X or Y chromosome number (in gonadal females: XO vs. XX, XX vs. XXY, XO vs. XY, XY vs. XXY; in gonadal males: XY vs. XXY, XY vs. XYY; XX vs. XXY, XO vs. XY).

**Results:** We measured numerous phenotypes to characterize these models, including gonadal histology, breeding performance, anogenital distance, levels of reproductive hormones, body and organ weights, and central nervous system sexual dimorphisms. Serum testosterone levels were comparable in adult XX and XY gonadal males. Phenotypes previously known to be sexually differentiated by the action of gonadal hormones were found to be similar in XX and XY rats with the same type of gonad, suggesting that XX and XY rats with the same type of gonad have comparable levels of gonadal hormones at various stages of development.

**Conclusion:** The results establish powerful new models to discriminate sex-chromosome and gonadal hormone effects that cause sexual differences in rat physiology and disease.

**Plain English Summary:** The Four Core Genotypes and XY* mouse models have been broadly useful for determining if sex differences in any mouse phenotype are caused by gonadal hormones, or by sex-chromosome complement (XX vs. XY), and if sex-chromosome effects are caused by X- or Y-linked mechanisms. Using gene knockout and transgenic methods, we have produced laboratory rat models that offer similar capabilities. The new rat models allow investigators to test with relative ease, for the first time, if a sex difference in a rat trait is caused by effects of XX vs. XY sex chromosomes, not mediated by effects of gonadal hormones, and to narrow the search for X or Y genes that have that role. The models produce XO, XX, XY, and XXY rats with ovaries, and XO, XX, XY, XXY, and XYY rats with testes. The four XX and XY groups represent a Four Core Genotypes rat model, comparison of which tests for sex-chromosome and gonadal hormonal effects that cause female and male rats to have different physiological or disease traits. Moreover, comparison of rats with different numbers of X chromosomes, or of Y chromosomes, but with the same type of gonad, provides evidence regarding the effects of X or Y dosage on rat traits. The new models will improve understanding of the impact of sex chromosomes on diseases or traits that are best modeled in rats. They will also improve understanding of the evolution of functional roles of sex chromosomes.

**Highlights:** It is advantageous to establish the factors that cause sex differences in diseases, because those factors mitigate or exacerbate diseases. We have produced new laboratory rats that have different types and numbers of sex chromosomes but the same type of gonad, allowing investigation of the role of sex chromosomes in causing sex differences in physiology and disease. The new rat lines allow comparison of XX and XY rats with the same type of gonad, to detect sex differences caused in part by the sex chromosomes. Other comparisons of rats with the same gonad but with different numbers of X chromosomes (XO vs. XX, XY vs XXY) or of Y chromosomes (XO vs. XY, XX vs. XXY, XY vs. XYY) detect effects of X or Y chromosome number. These resources can uncover sex-chromosome effects on any rat phenotype.

## Introduction

Biological sex differences in physiology and disease are caused by two main classes of sex-biased factors, gonadal hormones (ovarian vs. testicular), and sex-chromosome complement (XX vs. XY) [1]. Distinguishing the effects of sex-chromosome complement has been challenging, because the sex chromosomal and hormonal factors are normally tightly linked in comparisons of typical XX gonadal females and XY gonadal males. The Four Core Genotypes (FCG) mouse model has been useful in separating these effects, by unlinking sex-chromosome complement from differentiation of gonads [2–4]. In FCG mice, the testis-determining gene *Sry* is deleted from the Y chromosome (ChrY), producing a variant (“Y^−^“) that fails to cause testis development but allows ovarian differentiation [5]. The insertion of a functional *Sry* transgene onto an autosome complements the loss of *Sry*, so that XY^−^(*Sry*TG+) mice have testes and are fertile [6, 7]. When mated with XX females, they produce four types of offspring: XX and XY mice with the *Sry* transgene and testes, and XX and XY mice lacking *Sry* and possessing ovaries. In this model, sex-chromosome effects on any phenotype are measured by comparing XX and XY mice with the same type of gonad (twice, in mice with testes or ovaries), and hormone effects are measured by comparing mice with different types of gonads but with the same type of sex chromosomes (twice, in either XX or XY). Use of FCG mice has led to the novel discovery of sex-chromosome effects that contribute to sex differences in body weight and adiposity, autoimmune disease, Alzheimer’s disease, numerous cardiovascular diseases, neural tube closure defects, and the brain and behavior [8–16]. Use of FCG mice confirmed gonadal hormonal control of other sex differences [2, 14, 17, 18].

Once a sex-chromosome effect is detected in FCG mice, the XY* mouse model can be used to determine if the effect is attributable to variations in the number of X chromosomes (ChrX) or ChrY [4, 7]. The XY* model compares mice with one or two ChrX, keeping the number of ChrY constant (XO vs. XX gonadal females, XY vs. XXY gonadal males). The XY* model also compares mice with and without a ChrY (XO vs. XY, XX vs. XXY) to detect ChrY effects. The combined use of the FCG and/or XY* models has led to the discovery of several X and Y genes whose dose contributes to sex differences in diverse phenotypes [3, 8–10, 19].

A significant limitation is that the FCG and XY* models have only been available in mice. It is important to avoid an overreliance on a single species as the only animal model system in which important biomedical questions can be addressed. Any fundamental question of biology, such as the importance of sex-biasing effects of sex chromosomes in non-gonadal tissues, must be studied in diverse species to achieve a general answer. Here we report the development of laboratory rat (*Rattus norvegicus*) lines that offer some of the advantages of the FCG and XY* models, to detect sex-chromosome effects and dosage effects of ChrX and ChrY. Thus, the rat becomes the second mammalian species in which sex-chromosome effects can be studied with relative ease [20]. Rats have many advantages as a model species because of the vast literature on rat sex differences, their greater utility in some models of human disease, and their larger size [21–24]. Transgenesis is used increasingly in rats, and rat genomic and epigenomic resources are being rapidly developed [25–31].

We produced new rat lines by following the approach used to make FCG mice. We generated a rat ChrY that does not cause formation of testes, and inserted a functional *Sry* transgene into an autosome. A significant complication is that rats possess 8-11 different *Sry* genes on ChrY, depending on strain [32, 33]. The specific *Sry* gene(s) causing the differentiation of testes are not known. We previously reported that at least three *Sry* genes (*Sry4A*, *Sry1*, and *Sry3C*) are expressed in the gonadal ridge as it begins testis differentiation at embryonic day 13 (21 tail somites) in Sprague Dawley rats [34]. Therefore, as a transgene, we used a single Bacterial Artificial Chromosome (BAC) clone encoding all three of these *Sry* genes and no other genes. We report here the production of XY gonadal males with an autosomal *Sry* BAC transgene that causes testis development in XX offspring, allowing comparison of XX and XY gonadal males. To interfere with the testis-determining function of ChrY, we utilized CRISPR gRNAs aimed at ChrY regions near *Sry4A* and *Sry1*. This method produced fertile XY gonadal females that had XX and XY offspring with ovaries. Together, these resources provide an avenue for investigating sex-chromosome effects in virtually any rat phenotype. We report here the development of these resources and initial investigations of phenotypes.

## Methods

All experiments were approved by the Institutional Animal Care and Use Committees of participating institutions. Here, “male” and “female” are defined by type of gonad.

### Production of XX(SryTG+) gonadal male rats

Based on our previous evidence for expression of *Sry4A, Sry3C,* and *Sry1* in the embryonic gonadal ridge at the onset of rat testis development [34], we selected a single BAC clone RNECO-180E22 (Genbank AC279199) encoding these genes and no other known genes. At the Medical College of Wisconsin (MCW), we demonstrated by pronuclear microinjection into zygotes that a transgene, comprising this BAC clone and the pCC1BAC plasmid vector, caused formation of testes in a single transgenic XX female founder rat. Because XX gonadal males lack sperm, the model could not be propagated. At the University of Michigan Transgenic Animal Model Core, this BAC was purified and injected into pronuclei of more Sprague Dawley zygotes (Charles River Laboratories, Crl:CD(SD) strain code 001) to generate 22 founder rats carrying the transgene. Of 22 founders, 11 were XX, of which 6 were gonadal males. Four lines of XY transgenic founders [CD- Tg(RNECO-180E22)208Arno (RGD:155663365), CD-Tg(RNECO-180E22)733Arno (RGD:155663367), CD- Tg(RNECO-180E22)424Arno (RGD:155663366), and CD-Tg(RNECO-180E22)737Arno (RGD:155663368)] transmitted the transgene to XX gonadal male progeny. Hereafter we simply refer to the line numbers 208, 733, 424, and 737. We backcrossed all four transgenic lines to the Sprague Dawley Crl:CD(SD) strain code 001 (Charles River, Wilmington MA) for >10 generations to isolate a single transgene integration site in each line, and to measure phenotypes. Mating XXWT females to XY(*Sry*TG+) males produced XXWT gonadal females, and three types of gonadal males: XYWT, XX(*Sry*TG+) and XY(*Sry*TG+) (Figure 1, Tables 1 and 2, hereafter called XYWT, XXTG and XYTG, respectively). Offspring genotype was determined by PCR as discussed below (Table 3).

**Figure 1.**
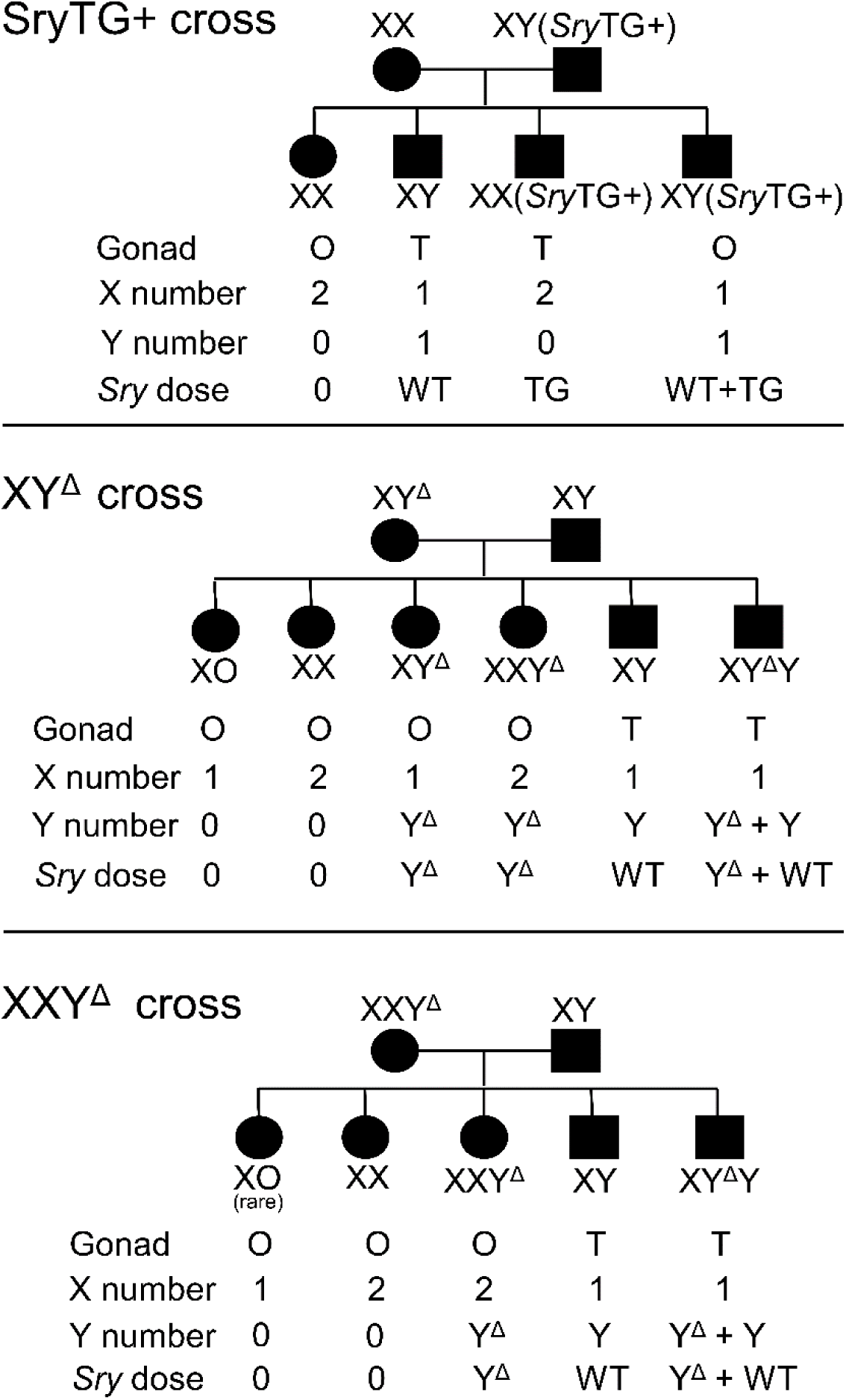
Three crosses producing sex-chromosome variant rats. Above: The *Sry* autosomal transgene is carried by the XY(*Sry*TG+) father, which produces four types of sperm, X and Y each with or without the transgene. Middle: Crossing XY^Δ^ females to XYWT males produces six types of viable genotypes, indicating substantial nondisjunction of X and Y chromosomes during meiosis in oogonia. Below: Crossing an XXY^Δ^ mother with XYWT father produces four types of progeny, indicating that XXY^Δ^ mothers typically produce X or XY^Δ^ eggs. One XXY^Δ^ mother produced a single rare XO offspring (Table 2). XY^Δ^ or XXY^Δ^ mothers may be bred with XY(*Sry*TG+) males, producing the illustrated offspring types with the addition of the *Sry* transgene. These crosses allow informative comparisons listed in Table 1.

**Table 1.**
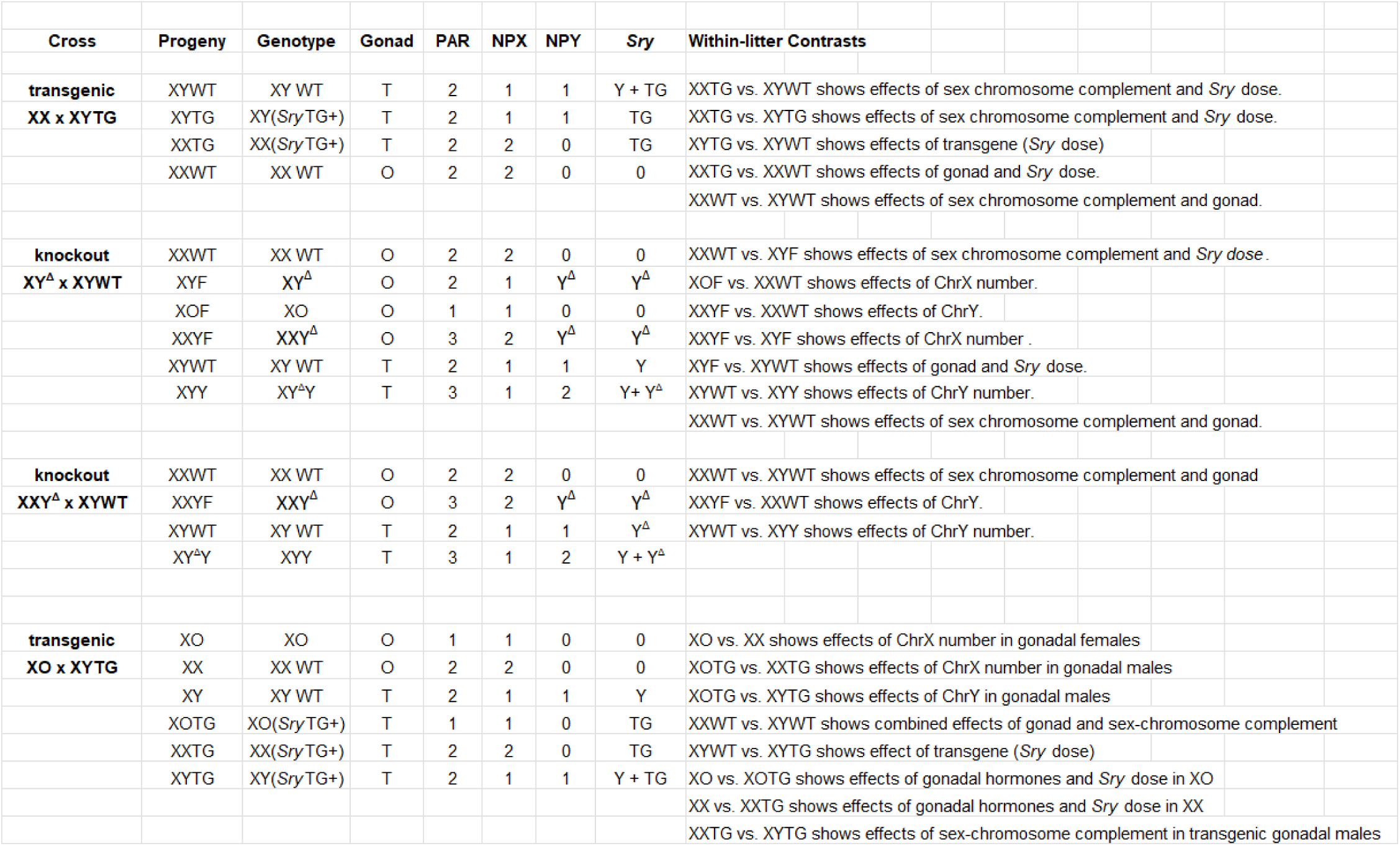
Genotypes produced in several breeding schemes, and contrasts for measuring effects of sex chromosome complement, gonad, ChrX number, and ChrY number.

**Table 2.**
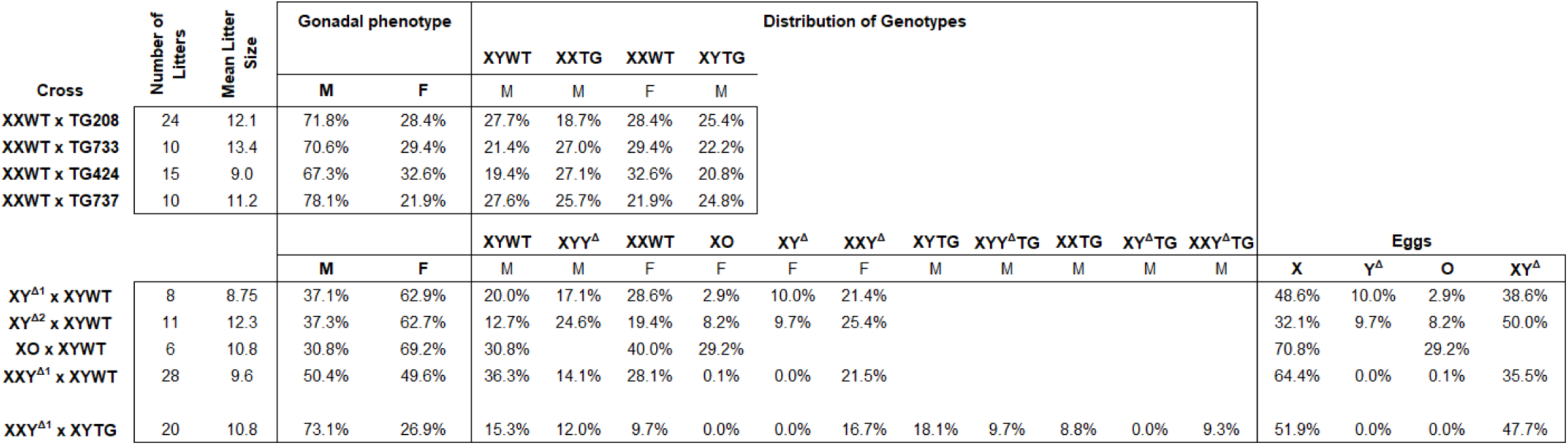
Frequencies of Genotypes.

**Table 3:**
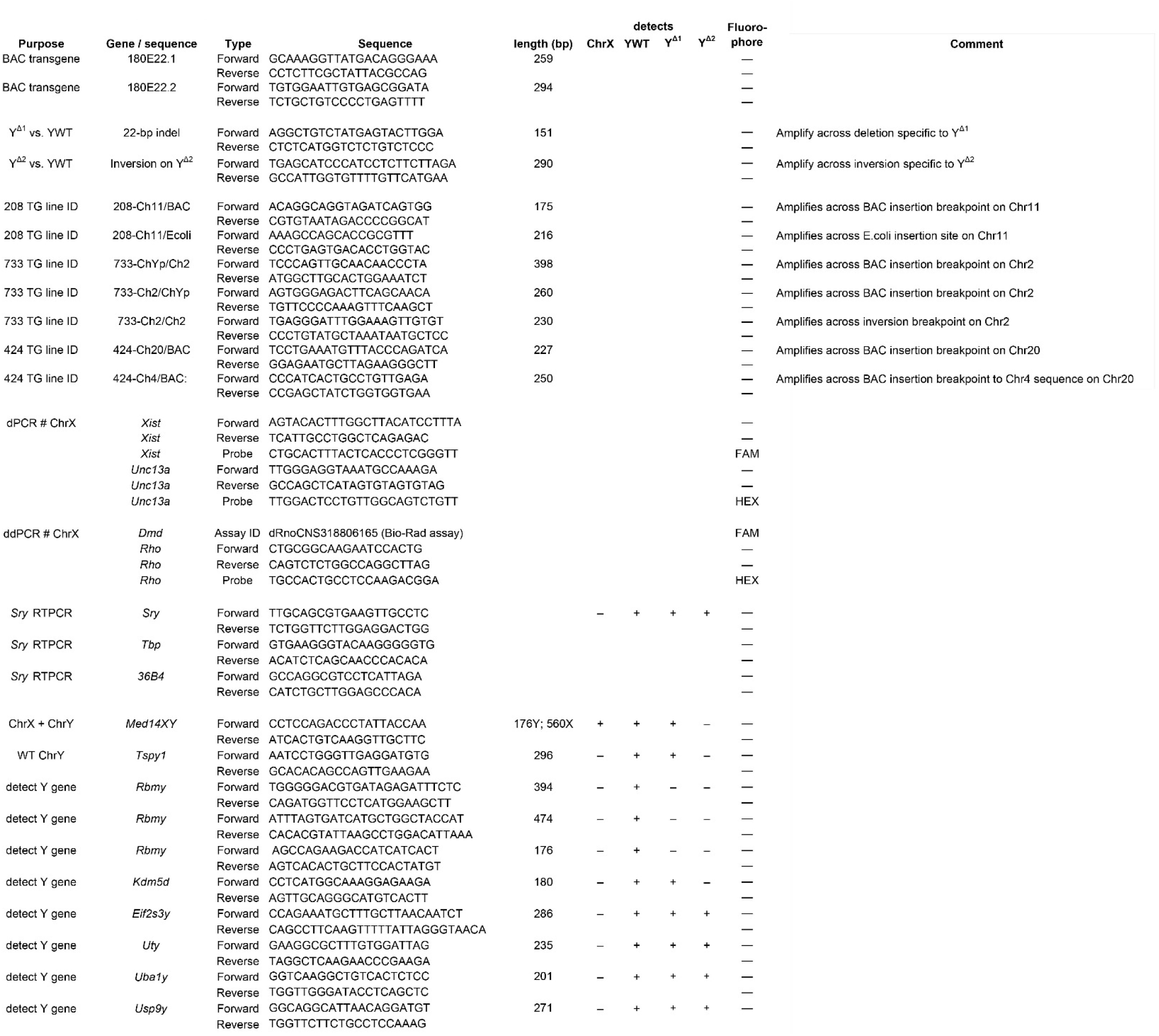
PCR primers and probes.

### Production and genomic characterization of two lines of XY***^Δ^*** gonadal female rats

The same BAC clone RNECO-180E22 sequence was used to design SpyCas9 guide RNAs flanking ∼67 kilobases of ChrY sequence harboring *Sry4A* and *Sry1* where unique sites could be identified in an attempt to delete both genes (Figure 2A). XY^Δ^ mutant rats [SD-Del(Yp)1Mcwi (RGD:155663364)] were generated at MCW by pronuclear injection of these two CRISPR-Cas9 ribonucleoproteins (RNPs) targeting the sequences GCATGTGGGCAGTTTCCACC<u>TGG</u> and ACACAGCTCCTCTCTGGTAG<u>AGG</u> (protospacer adjacent motif underlined) into single-cell Crl:SD (Charles River Laboratories, Crl:SD strain code 400) embryos. A single gonadal female carrying ChrY was identified, designated XY^Δ1^. This female was fertile and successfully transmitted the mutant ChrY following backcross to a wild-type XY Crl:SD male. Mating XY^Δ1^ females to XYWT males produced litters with females (XXWT females, XY^Δ1^, XXY^Δ1^, or XO) and males (XYWT or XYY^Δ1^). (Figure 1, Tables 1 and 2). Thus, XY^Δ1^ females produced four types of egg cells (“O”, X, Y, XY) because of nondisjunction at the first meiotic division [35]. XY^Δ1^ or XXY^Δ1^ females were backcrossed to Crl:SD and Crl:CD(SD) XYWT males for more than 19 generations.

**Figure 2:**
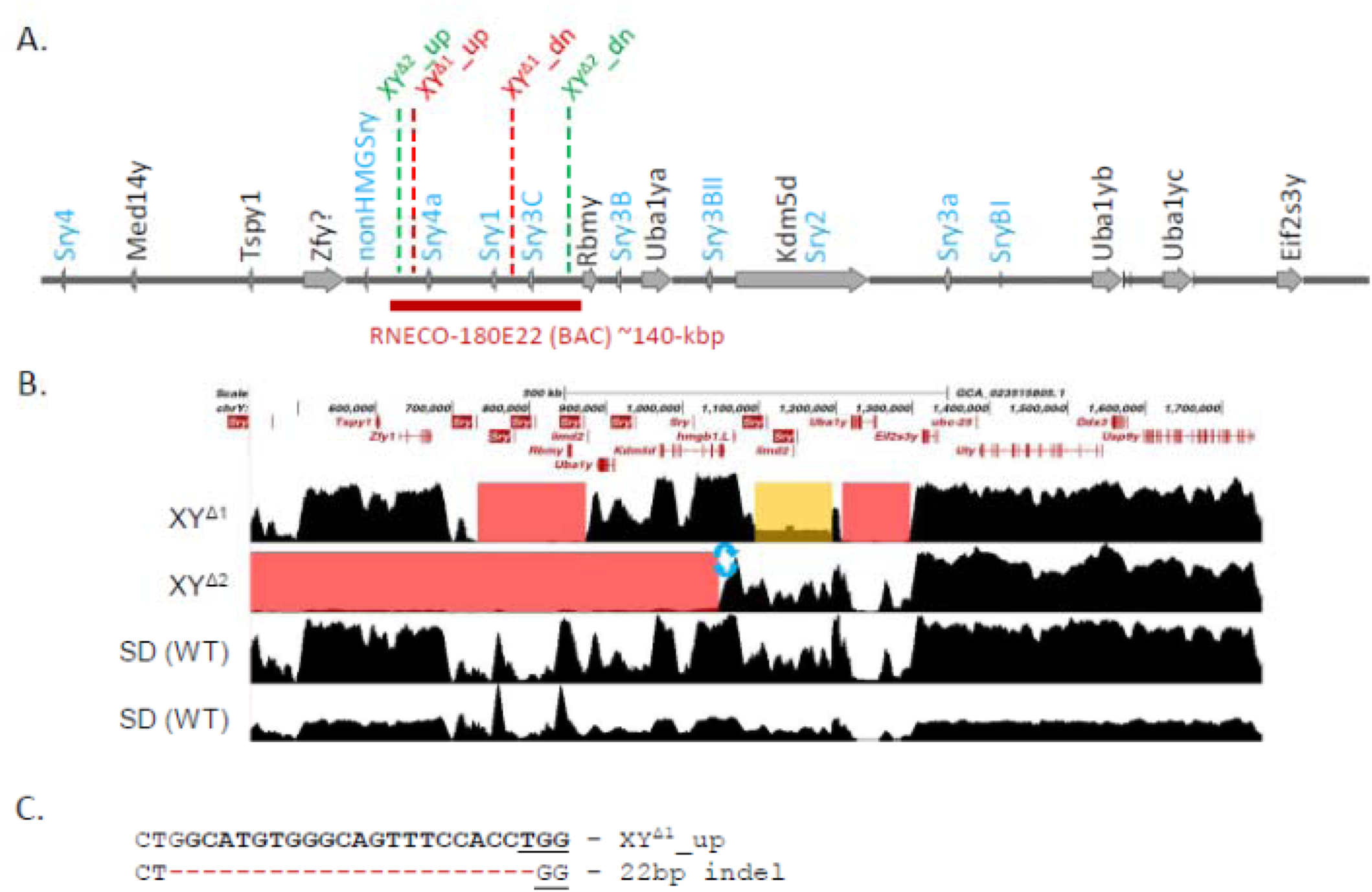
Genomic characterization of XY^Δ^ models. A. Schematic map showing the order of genes along a ∼1-Mbp region of the SHR/Akr rat Yp chromosome, based on Genbank accession OP905629. Ten Sry genes are shown. It is unknown if Sprague Dawley rats have an *Sry3* gene (GenBank: EU984077) in this or another region of ChrY. *Sry3* has thus far only reported in SHR strain, but not the Wistar-Kyoto (WKY) strain [31]. Positions of CRISPR target sites to generate the XY^Δ1^ and XY^Δ2^ are shown in red and green, respectively. The location of the RNECO-180E22 BAC sequence is shown. B) Among the currently available rat genome assemblies, OP905629 is most contiguous with the WKY/Bbb strain whole genome (assembly GCA_023515805.2, data not shown). The paired-end sequence coverage of ChrY in the XY^Δ1^ and XY^Δ2^ and two Sprague Dawley control animals aligned to the WKY/Bbb is displayed. Deletions are highlighted in red boxes in both lines and one putative deleted region in XY^Δ1^ in orange. *Rbmy* is deleted in both XY^Δ^ models. C) The XY^Δ1^ model also harbors a 22-bp deletion induced by gene editing where the XY^Δ1^_up guide RNA targets the chromosome and a small inversion near the 3’ end of *Kdm5d* was detected in XY^Δ2^ (blue arrows in panel B). Primers specific to these regions can be used to distinguish the mutant from wild-type ChrY by PCR (see Table 3 for details).

At early stages of breeding XY^Δ1^ females to XYWT males, we failed to discriminate XY^Δ1^ and XXY^Δ1^ offspring, and therefore inadvertently produced increasing numbers of fertile XXY^Δ1^ females. The XXY^Δ1^ females were bred based on the incorrect assumption that they were XY^Δ1^. This error resulted in the eventual loss of XY^Δ1^ females from our colonies, because XXY^Δ1^ mothers do not produce XY^Δ1^ daughters (Figure 1, Table 2). Some phenotypic analyses presented here were thus performed on a group composed of a mixture of XY^Δ1^ and XXY^Δ1^ genotypes. Digital PCR was used to resolve the genotypes of Y^Δ1^-bearing females after the fact. The separation into two groups of Y^Δ1^-bearing females (XY^Δ1^ and XXY^Δ1^) resulted in smaller group sizes for some phenotypic data collected from these females.

To recapture an XY^Δ^ female group, we produced a second line, XY^Δ2^, using an alternative strategy. CRISPR-Cas9 RNPs targeted sequences flanking *Sry4A and Sry3C*, using the target sequences CACGCAGTAGTCAGCTGCTC<u>TGG</u> and CAACATGGGACTGATGGAAC<u>AGG</u> injected into single-cell Sprague Dawley Crl:CD(SD) zygotes (Charles River Laboratories, strain code 001). Multiple gonadal females were recovered and one produced the XY^Δ2^ genotype following backcross to Crl:CD(SD) XYWT males. Litters contained females (XXWT, XY^Δ2^, XXY^Δ2^, or XO) and males (XYWT or XYY^Δ2^). (Figure 1, Tables 1 and 2). Careful measurement of ChrX copy number by droplet digital PCR ensured that XY^Δ2^ females were routinely backcrossed to Crl:CD(SD) XYWT males for more than 8 generations. When referring simply to XY^Δ^, we are referring to either the XY^Δ1^ or XY^Δ2^ models.

### Genome sequence analysis of transgenic and knockout lines

To provide a preliminary analysis of the CRISPR-induced genomic lesion in the XY^Δ1^ and XY^Δ2^ and the four transgenic (XYTG) lines, genomic DNA was harvested from spleen of individual mutant rats and two control Crl:SD animals and prepared for Illumina 150-bp paired-end whole genome sequencing (WGS) by the MCW Mellowes Center for Genomic Sciences and Precision Medicine on the Novaseq S4 flow cell to generate 30X coverage.

Quality-filtered reads were mapped to the WKY/Bbb genome (GCA_023515805.2) using a custom, lane-aware preprocessing pipeline incorporating Picard Tools v3.3.0 and BWA-MEM v0.7.17. The pipeline performed read group assignment, adapter marking, alignment, and merging of lane-specific BAMs to produce high-quality, duplicate-marked BAM files suitable for downstream breakpoint detection and variant analysis.

XXTG rats from lines 208, 424, and 733 were additionally sequenced using PacBio HiFi sequencing provided by Azenta Life Sciences to detect autosomal integration sites. Sequenced reads were mapped independently with minimap2 [36] to the Dahl SR_JrHsd strain (GCA_04568800 5.1). This assembly was chosen among the available inbred strain genome assemblies because it is phylogenetically most closely related strain to Sprague Dawley [37]. The Dahl_SR_JrHsd assembly was modified by removing the Y chromosome (CM099021.1). The DH10B laboratory strain *E. coli* genome (U00096.3), and the BAC vector sequence (AC279199.1) were also included in the reference genome. Once complete, the reference genome was indexed using the minimap2 index function. The reads were mapped and subsequently converted from SAM format to BAM format, coordinate sorted and again indexed using Samtools [36].

### Genotyping

Offspring genotype was determined by PCR. DNA was extracted from tails by using Chelex resin (Bio-Rad Laboratories, Hercules, CA), or by isopropanol precipitation from proteinase K-digested tissue. The presence of ChrY was detected using primers that amplify Med14X and Med14Y producing a 560 bp product for Med14X and a 172 bp product for Med14Y (Table 3). Presence of the *Sry* BAC transgene was determined using primers spanning each end of the 180E22 BAC clone and the adjacent pCC1BAC vector sequence with expected PCR product 259bp (primer pairs 180E22.1) or 294bp (primer pairs 180E22.2, Table 3). The Y^Δ1^ chromosome was detected by amplifying a unique sequence with expected PCR product of 218 bp compared to 239bp from intact ChrY (Table 3). The presence of the Y^Δ2^ chromosome was detected using primers amplifying a small inversion sequence of 290bp detected by NGS sequencing (Table 3). Other primers were used to detect unique sequences in each of the BAC transgenic lines, or were Y chromosome markers used to characterize the XY^Δ^ models (Table 3).

Digital PCR was used to determine the number of ChrX, using two methods. Method 1 (UCLA): PCR was performed using the QuantStudio Absolute Q system (Thermo Fisher Scientific, Los Angeles, California) (Table 3). Copy number of ChrX gene *Xist* was compared to the autosomal reference gene *Unc13a*. The *Xist* probe was detected using FAM fluorescence, and the *Unc13a* probe using *HEX* fluorescence. Absolute Q Digital PCR System software (Thermo Fisher Scientific) was used to determine thresholds and calculate target concentrations. Method 2: (Medical College of Wisconsin and Hudson Institute): droplet digital PCR was performed using the Bio-Rad QX200 droplet digital PCR system, including the QX200 droplet generator, C1000 Touch™ thermal cycler, and QX200 droplet reader. The ChrX gene *Dmd* was quantified using a Bio-Rad predesigned droplet digital PCR Copy Number Assay (Rat Unique Assay ID: dRnoCNS318806165). The autosomal control gene *Rho* was measured using a custom HEX-labeled probe (Table 3). Thermocycling conditions were 95°C for 10 minutes, followed by 40 cycles of 94°C for 30 seconds and 60°C for 1 minute, with a final enzyme deactivation step at 98°C for 10 minutes. QuantaSoft Software (Bio-Rad) was used to determine thresholds and calculate target concentrations.

### Fluorescent *in situ* hybridization (FISH) localization of Sry transgene in metaphase chromosomes

Preparation of metaphase spreads from skin fibroblasts, fluorescent labeling of the *Sry* BAC clone RNECO-180E22, and hybridization of the probe to metaphase spreads have previously been described [38]. Skin fibroblasts were prepared by mincing ear biopsies and culturing in Advanced Dulbecco’s Modified Eagle’s Medium and antibiotics. Metaphase chromosome spreads were prepared by inhibiting cell division with colcemid, followed by cell dispersion, centrifugation, and treatment with KCl and 3:1 methanol:acetic acid. The rat BAC clone RNECO-180E22, encoding *Sry4A*, *Sry1*, and *Sry3C* was fluorescently labeled using the Invitrogen (Carlsbad, California) FISH Tag_TM_ DNA Kit. The probe was hybridized to metaphase spreads at 37°C for 24 hours, washed, dehydrated, dried, counterstained with DAPI, and viewed under a microscope.

### Liver transcriptome analysis

To determine if inserts of *E. coli* genome sequences into transgenic rats resulted in expression of *E. coli* transcripts, paired-end RNAseq was performed on RNA extracted from livers of one XYWT male, and one XYTG male each from lines 208 and 733 (Azenta Life Sciences, Burlington MA), with 26 to 31 million reads per sample. Reads were assessed for multiple species using the PlusPFP library with KRAKEN2 analyses [39].

Reads were then concatenated from all samples and built into a *de novo* Trinity assembly [40] using Trimmomatic read cleaning [41], followed by Transdecoder and Trinotate annotation of gene and species matching [42].

### Gonadal histology

Testes and ovaries were collected at approximately postnatal day 60 (∼P60, P1 is the day of birth) and immersed in freshly prepared 4% paraformaldehyde (PFA) overnight at 4°C. They were transferred to 70% alcohol before embedding in paraffin for later sectioning. The gonads were sectioned at 4µm and sections collected on Superfrost slides. Hematoxylin and Eosin staining was performed by the Monash Histology Facility using established protocols [43]. Stained slides were imaged using the Leica Aperio VS120 Microscope and processed using the Leica ImageScope software.

### *Sry* expression in tissues

Spleen, liver, and gonads were collected from P60-65 male rats from lines 208 and 733 at backcross generation 7, and from XX and XY^Δ1^ female rats at backcross generation 4. They were flash frozen in liquid nitrogen, then stored at −80°C until further use. RNA was isolated from tissues with Trizol (Invitrogen, Carlsbad, CA, USA) and treated with ezDNase (Invitrogen) to prevent possible contamination with genomic DNA. First-strand cDNA was generated using cDNA reverse transcription kit (Applied Biosystems, Foster City, CA, USA) using oligo dT12-18 primers (Invitrogen). Quantitative real time PCR (n□=□7–8 per genotype) was performed on an ABI 7300 Sequence Detection system (Applied Biosystems) using the PowerUp SYBR Green Master Mix Kit (Applied Biosystems).

Pan-*Sry* primers were used to amplify all the *Sry* genes except the *nonHMGSry* gene. TATA box-binding protein and 36B4 were used as loading control genes (Table 3). Cycling conditions were: 95°C for 3 min; 40 cycles of 95°C for 15 sec, 60°C for 30 sec and 72°C for 30 sec with standard curve for *Sry* gene and control genes with four serial dilution points of control cDNA. Dissociation curves were examined to eliminate the possibility of genomic DNA contamination. Expression of *Sry* was expressed as fold change relative to the control gene.

### Growth curves and organ weights

Rats were group-housed and maintained at 23°C on a 12:12 light/dark cycle. Rats were fed regular chow diet with 5% fat (LabDiet 5001, St. Louis, MO, USA). Body weight was measured at P21 and then at weekly intervals until ∼P60, at backcross generation 4 for females and generation 7 for males. At ∼P60, spleens, kidneys and gonads were dissected and weighed.

### Anogenital distance

Anogenital distance (AGD) was measured with calipers at P6-7 at backcross generation 7 for transgenic litters. It was also measured in progeny of the cross XXY^Δ1^ x XYTG at backcross generation 9 or 11. Because of variability in body size across litters, each rat’s AGD was expressed as a ratio relative to the mean AGD of XYWT rats in its litter.

### Serum testosterone, follicle-stimulating hormone (FSH) and luteinizing hormone (LH) levels

Whole blood was collected using cardiac puncture from isoflurane-anesthetized rats at ∼P60. Blood was allowed to clot at room temperature for >60 minutes and then centrifuged at 1000g for 10 min to collect serum, then stored at -80°C until analyzed. At backcross generation 4, serum from transgenic lines 208 and 733 was analyzed for testosterone or FSH or LH by the Ligand Assay and Analysis Core at the University of Virginia. At background generation 7, serum from all transgenic lines was analyzed by a competitive immunoassay using the Testosterone Elisa Kit (Enzo Life Sciences Inc., Farmingdale, New York) with sensitivity at 5.67pg/ml and range 7.81 - 2,000 pg/ml. The data were analyzed by using a Four Parameter Logistic Curve from MyAssays.

### Histological analysis of sexually dimorphic spinal nuclei

Adult P59-63 rats at backcross generation 4 from line 208 were anesthetized with isoflurane and perfused with heparinized phosphate buffered saline, and then with 4% paraformaldehyde (PFA). Spinal cords from a total of 48 animals (XYWT, n = 10; XYTG, n = 10; XXTG, n = 9; XY^Δ^, n = 2; XXY^Δ^, n = 4; XXWT, n = 13) were embedded in gelatin, frozen, and sectioned transversely at 40 µm. Sections were mounted on gelatin-coated slides and stained with thionin.

### Number of Motoneurons

The rat Spinal Nucleus of the Bulbocavernosus (SNB) and Dorsolateral Nucleus (DLN) are discrete groups of motoneurons located in the L5/L6 spinal segments [44–46], which innervate perineal muscles involved in male sexual function. Sex differences in the number and size of motoneurons in these nuclei are known to be caused by sex differences in the secretion of testosterone perinatally [47]. For each animal, the total numbers of SNB and DLN motoneurons were estimated using the optical disector method (Stereo Investigator, MBF Bioscience, Williston, VT). Counts were made at x937.5 under brightfield illumination. SNB and DLN motoneurons were easily recognizable as large, darkly staining, multipolar cells. In the SNB, a counting frame (110 µm X 80 µm) was moved systematically throughout a midline area (∼500 µm x ∼300 µm) just ventral to the central canal. For the DLN, the counting frame (∼500 µm x ∼500 µm) was moved along the lateral margin of the ventral horn. Only motoneurons in which there was a clear nucleus and nucleolus were counted, provided that they did not contact the forbidden lines of the counting frame.

Motoneuron nucleoli were counted as they appeared while focusing through the z-axis, and nucleoli in the first focal plane (i.e., “tops”) were excluded to avoid double counting. The length of the disector was approximately 16 µm, which was adequate for visualizing nucleoli in multiple focal planes. Motoneuron counts in the SNB were derived from a mean of 16.1 sections spaced 160 µm apart and distributed uniformly through the entire rostrocaudal extent of the SNB, or in the DLN from a mean of 15.6 sections spaced 160 µm apart. This sampling scheme produced an average of estimated coefficient of error (CE) of 0.09 for the SNB and 0.07 for the DLN. Cell counts for each animal were corrected for the proportion of sections sampled.

### Soma size

The cross-sectional area of SNB and DLN motoneuronal somata was assessed using the Nucleator method with Stereo Investigator. A set of four rays emanating from a point randomly chosen within each motoneuron soma was drawn and oriented randomly. Soma areas of an average of 41.7 SNB and 22.9 DLN motoneurons were measured for each animal at a final magnification of x780 under brightfield illumination; average estimated coefficient of error (CE) was 0.01 for SNB and 0.02 for DLN. Soma areas within each animal were then averaged for statistical analysis.

### Histological analysis of limbic brain regions

#### Immunohistochemistry of the Sexually Dimorphic Nucleus of the Preoptic Area (SDN-POA)

The Sexually Dimorphic Nucleus of the Preoptic Area (a.k.a. caudal Medial Preoptic Nucleus) of the rat hypothalamus is a classic sexual dimorphism in a brain region controlling sexual function [48]. The sex difference is caused by perinatal sex difference in the levels of gonadal hormones [49]. Adult rats from line 208 at backcross generation 4 were anesthetized with isoflurane at P60-65 and perfused with saline and the 4% paraformaldehyde. Brains were removed, post-fixed with 4% paraformaldehyde (PFA), sectioned transversely at 40 μm. Sections were incubated in a primary antibody cocktail solution directed against calbindin (CALB, Mouse Monoclonal Anti-Calbindin-D-28K (Cat #C9848, Sigma, St. Louis, MO)) with a 1:10,000 dilution in 2% normal horse serum, 0.4% Triton X-100, and TBS for 48-72 hours at room temperature. Sections were then washed and placed in a solution of secondary antibody, biotinylated horse anti-mouse immunoglobulin G (IgG) (1:500, Vector Labs, Burlingame, CA), for 60 minutes at room temperature. The signal was developed using an avidin-biotin complex kit (Vector Labs, Burlingame, CA) and enhanced using DAB/cobalt chromagen (Sigma).

#### Immunohistochemistry of the Anteroventral Periventricular Nucleus (AVPV)

The AVPV is a sexually dimorphic region of the rat hypothalamus controlling ovulation, which is larger in females because of the effects of testicular hormones [50]. Adult rats of line 208 and progeny of XY^Δ^ females were studied at P60-65 at backcross generation 8 for males and 4 for females. Perfusion and sectioning of brains were as described for the SDN-POA. The sections were incubated for 30 minutes in a blocking solution, then incubated in the primary antibody cocktail for 48-72 hours (1:1000 dilution of tyrosine hydroxylase mouse monoclonal antibody, MAB318, Millipore Sigma). The secondary antibody was biotinylated horse anti-mouse, Vector Laboratories Kit PK-4002. Signal was developed using avidin-biotin complex enhanced with DAB/cobalt.

#### Quantification of calbindin-immunoreactivity (CALB-ir) in the SDN-POA

SDN-POA cell number, cell size, and nucleus volume was measured at 630x magnification. The anterior commissure marked the rostral extent of the SDN-POA, and its caudal extent coincided with the onset of CALB-ir cells in the Bed Nucleus of the Stria Terminalis [51]. CALB-ir cells within the SDN-POA region were found in 5-6 sections in males and 2-3 sections in females.

Images of brain sections were captured on a Leica DMRXA scope (Leica Microsystems, Wetzlar, Germany). The volume of the CALB-ir region of the SDN-POA was quantified by outlining the CALB-ir-defined nucleus and calculating the area of the nucleus on both sides of the brain in each section with ImageJ, then multiplying by section thickness (40 μm) and summing volumes of all sections. The number of CALB-ir cells was estimated using the optical disector method counting CALB-ir cells found within all sections through the nucleus, focusing through the Z axis and ignoring “tops” to avoid double counting. The total cell number was estimated as a sum of the counts collected from both hemispheres across all sections containing the SDN-POA. Mean cell size was estimated by measuring the areas of five randomly selected cells in each hemisphere in each section, resulting in 50-60 cell areas collected from the male SDN-POA and 20-30 cell areas measured from the female SDN-POA. The mean cell size was calculated for each animal from all the cells that were measured.

#### Quantification of tyrosine hydroxylase-immunoreactivity (TH-ir) in the AVPV

The AVPV area was visualized under a Leica brightfield microscope at 400x magnification. We measured 6 consecutive 40 µm sections, oriented by the morphology of the anterior commissure. The third section in the rostro-caudal series was the section at which the anterior commissure (AC) met at the midline [52]. TH-ir neurons were counted in two fields, each 100 by 200 μm. The ventral edges of the fields were at the junction of the third ventricle with the optic chiasm with the dorsal edge 200 μm above, and the medial edges of the fields were at the lateral edge of the third ventricle with field’s lateral edge 100 μm to the side. TH-ir neurons were counted using an optical disector method, excluding neurons at the top of sections to avoid double counting.

### Statistical analyses

For most measurements, experimental groups were compared using a one-way analysis of variance (ANOVA) across 3-6 genotypes (Figure 1) with a post hoc Tukey-Kramer (TK) test for pairwise comparisons (NCSS statistical software). For body weight measurements across ages, we used a two-way repeated measures ANOVA with a between-factor of genotypes and repeated factor of age to study the body growth curve in 3 male groups (from lines 208, 733, 424, and 737), and 3 female groups (XXWT, XY^Δ1^, and XXY^Δ1^) from XY^Δ1^ litters. Chi-square tests (χ^2^) were used to judge distributions of genotypes. In all figures, error bars show standard error of the mean (SEM). The results of statistical tests are in Table 4.

**Table 4.**
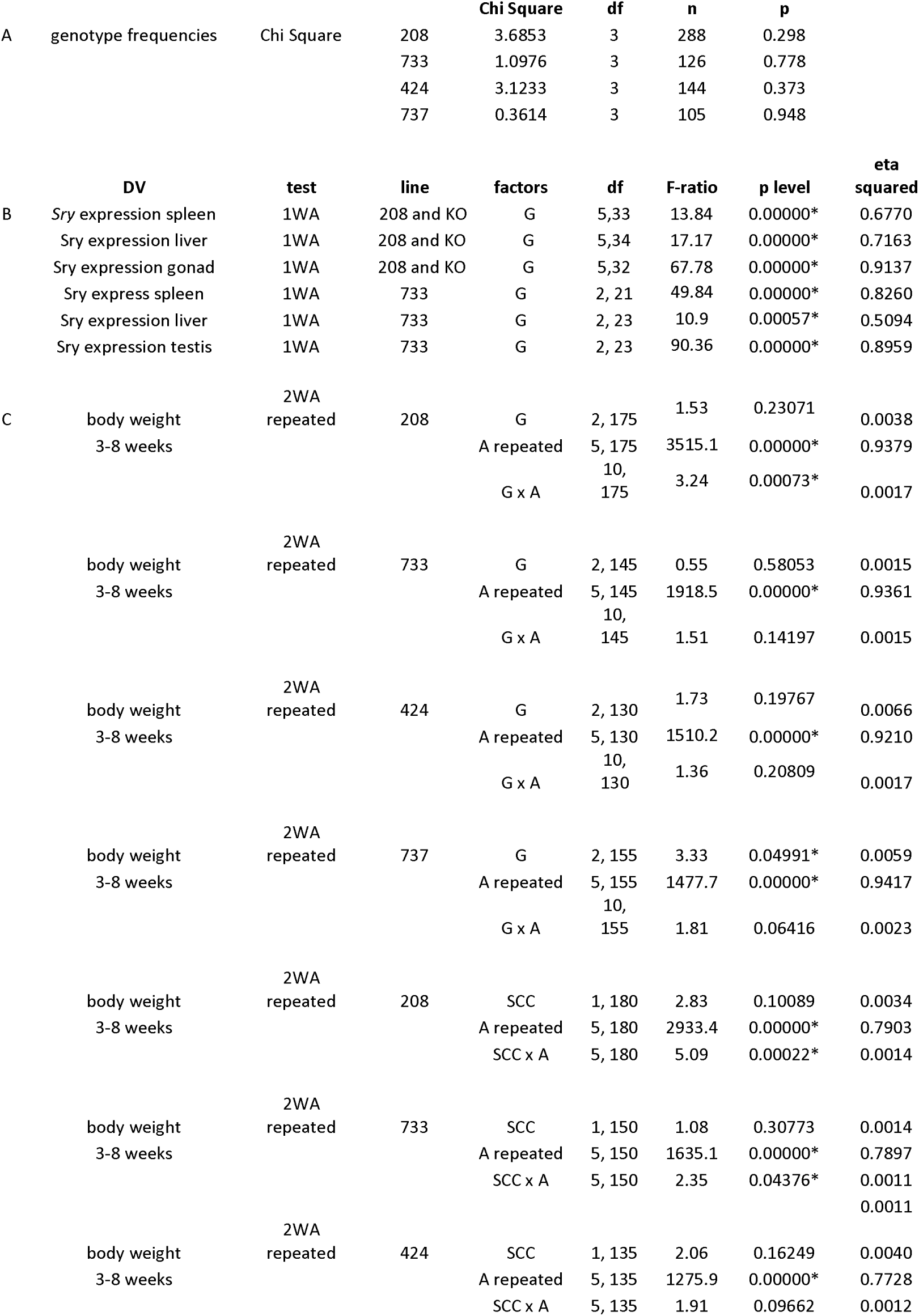
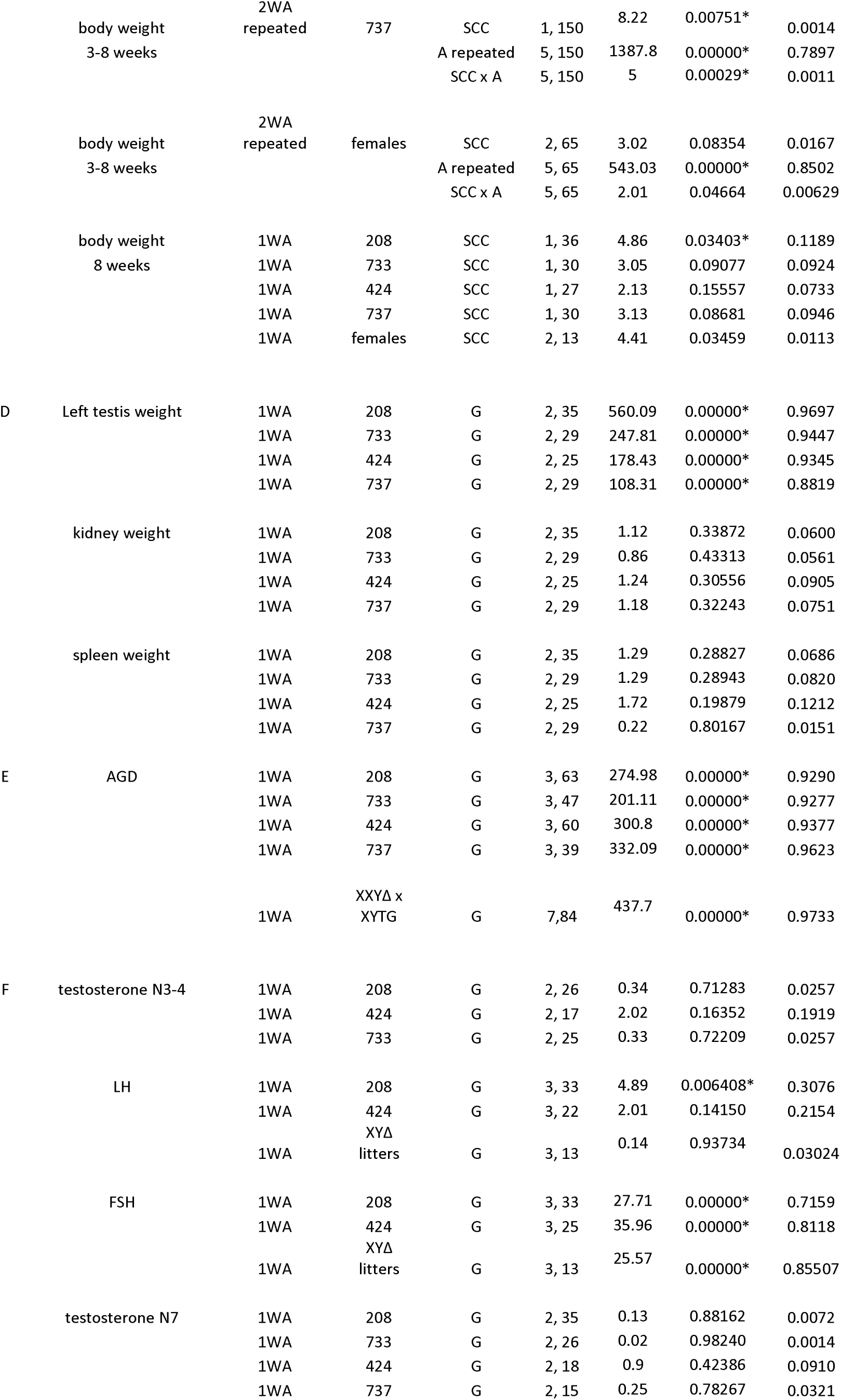
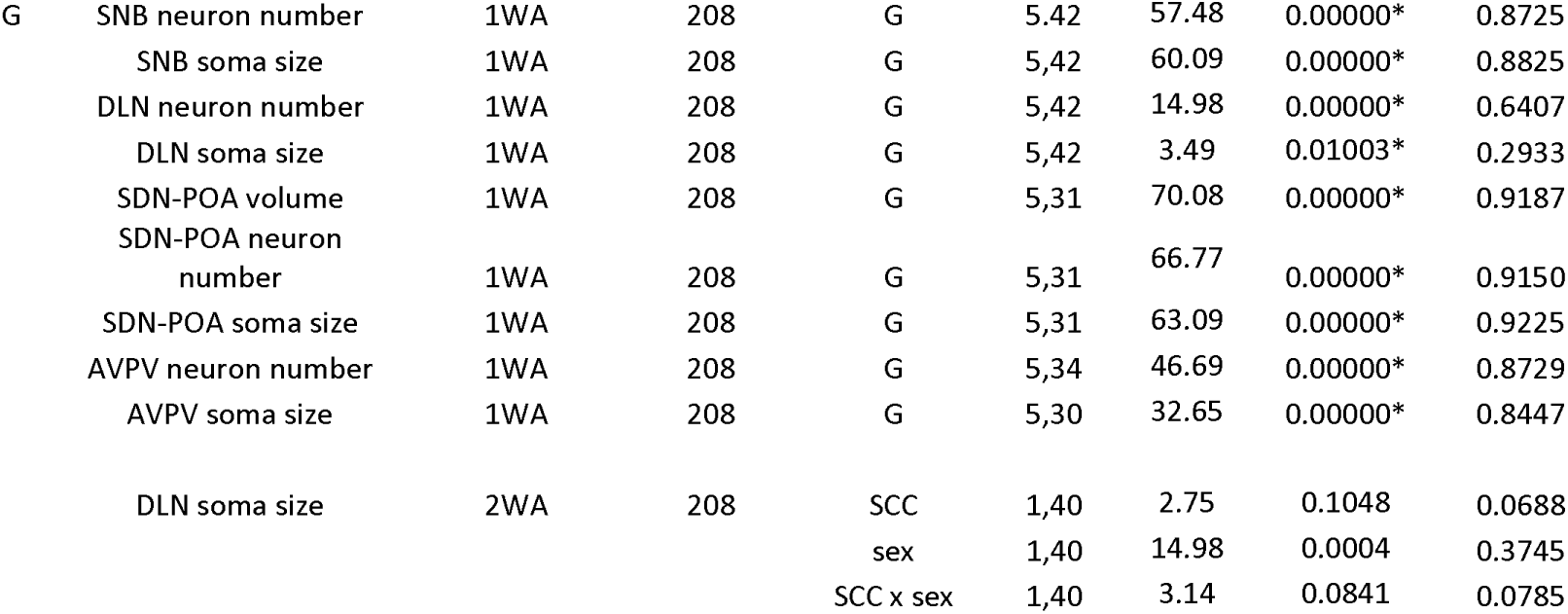
Results of statistical tests. Abbreviations: DV, dependent variable; 2WA, two-way ANOVA; 1WA, one-way ANOVA; A, age; G, genotype; SCC, sex chromosome complement (XX vs. XY, 2 levels); df, degrees of freedom; *significant at p<0.05 or better.

|  |  |  |  | Chi Square | df | n | p |  |
| --- | --- | --- | --- | --- | --- | --- | --- | --- |
| A | genotype frequencies | Chi Square | 208 | 3.6853 | 3 | 288 | 0.298 |  |
|  |  |  | 733 | 1.0976 | 3 | 126 | 0.778 |  |
|  |  |  | 424 | 3.1233 | 3 | 144 | 0.373 |  |
|  |  |  | 737 | 0.3614 | 3 | 105 | 0.948 |  |
|  | DV | test | line | factors | df | F-ratio | p level | eta squared |
| B | Sry expression spleen | 1WA | 208 and KO | G | 5, 33 | 13.84 | 0.00000* | 0.6770 |
|  | Sry expression liver | 1WA | 208 and KO | G | 5, 34 | 17.17 | 0.00000* | 0.7163 |
|  | Sry expression gonad | 1WA | 208 and KO | G | 5, 32 | 67.78 | 0.00000* | 0.9137 |
|  | Sry express spleen | 1WA | 733 | G | 2, 21 | 49.84 | 0.00000* | 0.8260 |
|  | Sry expression liver | 1WA | 733 | G | 2, 23 | 10.9 | 0.00057* | 0.5094 |
|  | Sry expression testis | 1WA | 733 | G | 2, 23 | 90.36 | 0.00000* | 0.8959 |
| C | body weight<br>3-8 weeks | 2WA<br>repeated | 208 | G | 2, 175 | 1.53 | 0.23071 | 0.0038 |
|  |  |  |  | A repeated | 5, 175 | 3515.1 | 0.00000* | 0.9379 |
|  |  |  |  | G x A | 10, 175 | 3.24 | 0.00073* | 0.0017 |
|  | body weight<br>3-8 weeks | 2WA<br>repeated | 733 | G | 2, 145 | 0.55 | 0.58053 | 0.0015 |
|  |  |  |  | A repeated | 5, 145 | 1918.5 | 0.00000* | 0.9361 |
|  |  |  |  | G x A | 10, 145 | 1.51 | 0.14197 | 0.0015 |
|  | body weight<br>3-8 weeks | 2WA<br>repeated | 424 | G | 2, 130 | 1.73 | 0.19767 | 0.0066 |
|  |  |  |  | A repeated | 5, 130 | 1510.2 | 0.00000* | 0.9210 |
|  |  |  |  | G x A | 10, 130 | 1.36 | 0.20809 | 0.0017 |
|  | body weight<br>3-8 weeks | 2WA<br>repeated | 737 | G | 2, 155 | 3.33 | 0.04991* | 0.0059 |
|  |  |  |  | A repeated | 5, 155 | 1477.7 | 0.00000* | 0.9417 |
|  |  |  |  | G x A | 10, 155 | 1.81 | 0.06416 | 0.0023 |
|  | body weight<br>3-8 weeks | 2WA<br>repeated | 208 | SCC | 1, 180 | 2.83 | 0.10089 | 0.0034 |
|  |  |  |  | A repeated | 5, 180 | 2933.4 | 0.00000* | 0.7903 |
|  |  |  |  | SCC x A | 5, 180 | 5.09 | 0.00022* | 0.0014 |
|  | body weight<br>3-8 weeks | 2WA<br>repeated | 733 | SCC | 1, 150 | 1.08 | 0.30773 | 0.0014 |
|  |  |  |  | A repeated | 5, 150 | 1635.1 | 0.00000* | 0.7897 |
|  |  |  |  | SCC x A | 5, 150 | 2.35 | 0.04376* | 0.0011 |
|  | body weight<br>3-8 weeks | 2WA<br>repeated | 424 | SCC | 1, 135 | 2.06 | 0.16249 | 0.0040 |
|  |  |  |  | A repeated | 5, 135 | 1275.9 | 0.00000* | 0.7728 |
|  |  |  |  | SCC x A | 5, 135 | 1.91 | 0.09662 | 0.0012 |
|  | body weight<br>3-8 weeks | 2WA<br>repeated | 737 | SCC | 1, 150 | 8.22 | 0.00751* | 0.0014 |
|  |  |  |  | A repeated | 5, 150 | 1387.8 | 0.00000* | 0.7897 |
|  |  |  |  | SCC x A | 5, 150 | 5 | 0.00029* | 0.0011 |
|  | body weight<br>3-8 weeks | 2WA<br>repeated | females | SCC | 2, 65 | 3.02 | 0.08354 | 0.0167 |
|  |  |  |  | A repeated | 5, 65 | 543.03 | 0.00000* | 0.8502 |
|  |  |  |  | SCC x A | 5, 65 | 2.01 | 0.04664 | 0.00629 |
|  | body weight<br>8 weeks | 1WA | 208 | SCC | 1, 36 | 4.86 | 0.03403* | 0.1189 |
|  |  | 1WA | 733 | SCC | 1, 30 | 3.05 | 0.09077 | 0.0924 |
|  |  | 1WA | 424 | SCC | 1, 27 | 2.13 | 0.15557 | 0.0733 |
|  |  | 1WA | 737 | SCC | 1, 30 | 3.13 | 0.08681 | 0.0946 |
|  |  | 1WA | females | SCC | 2, 13 | 4.41 | 0.03459 | 0.0113 |
| D | Left testis weight | 1WA | 208 | G | 2, 35 | 560.09 | 0.00000* | 0.9697 |
|  |  | 1WA | 733 | G | 2, 29 | 247.81 | 0.00000* | 0.9447 |
|  |  | 1WA | 424 | G | 2, 25 | 178.43 | 0.00000* | 0.9345 |
|  |  | 1WA | 737 | G | 2, 29 | 108.31 | 0.00000* | 0.8819 |
|  | kidney weight | 1WA | 208 | G | 2, 35 | 1.12 | 0.33872 | 0.0600 |
|  |  | 1WA | 733 | G | 2, 29 | 0.86 | 0.43313 | 0.0561 |
|  |  | 1WA | 424 | G | 2, 25 | 1.24 | 0.30556 | 0.0905 |
|  |  | 1WA | 737 | G | 2, 29 | 1.18 | 0.32243 | 0.0751 |
|  | spleen weight | 1WA | 208 | G | 2, 35 | 1.29 | 0.28827 | 0.0686 |
|  |  | 1WA | 733 | G | 2, 29 | 1.29 | 0.28943 | 0.0820 |
|  |  | 1WA | 424 | G | 2, 25 | 1.72 | 0.19879 | 0.1212 |
|  |  | 1WA | 737 | G | 2, 29 | 0.22 | 0.80167 | 0.0151 |
| E | AGD | 1WA | 208 | G | 3, 63 | 274.98 | 0.00000* | 0.9290 |
|  |  | 1WA | 733 | G | 3, 47 | 201.11 | 0.00000* | 0.9277 |
|  |  | 1WA | 424 | G | 3, 60 | 300.8 | 0.00000* | 0.9377 |
|  |  | 1WA | 737 | G | 3, 39 | 332.09 | 0.00000* | 0.9623 |
|  |  | 1WA | XXYΔ x<br>XYTG | G | 7,84 | 437.7 | 0.00000* | 0.9733 |
| F | testosterone N3-4 | 1WA | 208 | G | 2, 26 | 0.34 | 0.71283 | 0.0257 |
|  |  | 1WA | 424 | G | 2, 17 | 2.02 | 0.16352 | 0.1919 |
|  |  | 1WA | 733 | G | 2, 25 | 0.33 | 0.72209 | 0.0257 |
|  | LH | 1WA | 208 | G | 3, 33 | 4.89 | 0.006408* | 0.3076 |
|  |  | 1WA | 424 | G | 3, 22 | 2.01 | 0.14150 | 0.2154 |
|  |  | 1WA | XYΔ<br>litters | G | 3, 13 | 0.14 | 0.93734 | 0.03024 |
|  | FSH | 1WA | 208 | G | 3, 33 | 27.71 | 0.00000* | 0.7159 |
|  |  | 1WA | 424 | G | 3, 25 | 35.96 | 0.00000* | 0.8118 |
|  |  | 1WA | XYΔ<br>litters | G | 3, 13 | 25.57 | 0.00000* | 0.85507 |
|  | testosterone N7 | 1WA | 208 | G | 2, 35 | 0.13 | 0.88162 | 0.0072 |
|  |  | 1WA | 733 | G | 2, 26 | 0.02 | 0.98240 | 0.0014 |
|  |  | 1WA | 424 | G | 2, 18 | 0.9 | 0.42386 | 0.0910 |
|  |  | 1WA | 737 | G | 2, 15 | 0.25 | 0.78267 | 0.0321 |
| G | SNB neuron number | 1WA | 208 | G | 5,42 | 57.48 | 0.00000* | 0.8725 |
|  | SNB soma size | 1WA | 208 | G | 5,42 | 60.09 | 0.00000* | 0.8825 |
|  | DLN neuron number | 1WA | 208 | G | 5,42 | 14.98 | 0.00000* | 0.6407 |
|  | DLN soma size | 1WA | 208 | G | 5,42 | 3.49 | 0.01003* | 0.2933 |
|  | SDN-POA volume | 1WA | 208 | G | 5,31 | 70.08 | 0.00000* | 0.9187 |
|  | SDN-POA neuron number | 1WA | 208 | G | 5,31 | 66.77 | 0.00000* | 0.9150 |
|  | SDN-POA soma size | 1WA | 208 | G | 5,31 | 63.09 | 0.00000* | 0.9225 |
|  | AVPV neuron number | 1WA | 208 | G | 5,34 | 46.69 | 0.00000* | 0.8729 |
|  | AVPV soma size | 1WA | 208 | G | 5,30 | 32.65 | 0.00000* | 0.8447 |
|  | DLN soma size | 2WA | 208 | SCC | 1,40 | 2.75 | 0.1048 | 0.0688 |
|  |  |  |  | sex | 1,40 | 14.98 | 0.0004 | 0.3745 |
|  |  |  |  | SCC x sex | 1,40 | 3.14 | 0.0841 | 0.0785 |

## Results

### Production of XY***^Δ^*** lines

Figure 2A shows an annotated region of about 1.5-Mbp of assembled sequence (GenBank accession OP905629; SHR/Ark strain origin) harboring 10 *Sry* genes, other ChrY genes, as well as the positions of CRISPR guide RNAs used to generate female XY^Δ^ lines. It is not clear whether SD rats used in our studies harbor the *Sry3* gene, which would presumably be located outside this region.

Using two unique gRNAs to target sites flanking the region of *Sry4A* and *Sry1*, we produced a mutant founder XY^Δ1^ rat with ovaries that was fertile and produced both XXWT and XY^Δ1^ daughters (Table 2). A PCR amplicon spanning the XYΔ1_up target site amplified a product which harbors a 22-bp deletion at this site (Figure 2B) which can be used to distinguish the mutant ChrY ^Δ1^ from wildtype ChrY by PCR (data not shown). Paired-end Illumina sequence characterization of this model revealed unexpected outcomes on ChrY resulting from CRISPR-Cas9 mutagenesis when compared to the WKY/Bbb genome, which is more contiguous with the OP905629 sequence (data not shown). Multiple large segments are deleted in this model were evident as low or absent read coverage compared to two control SD animals (Figure 2B). Likely, the CRISPR-Cas9 RNPs cleaved ChrY at multiple sequences highly similar to the target site, resulting in multiple deletions and rearrangements of the chromosome. The deleted regions minimally include several *Sry* paralogs and *Rbmy*.

PCR amplification of genomic DNA from XY^Δ1^ rats, using unique primer sets within *Med14y*, *Tspy1*, *nonHMGSry*, *Uba1y, Kdm5d, Eif2s3y, Uty, and Usp9y,* confirmed that these genes are present (see Table 3). Thus, most or all of the broadly expressed Y genes [53] are present.

A second model XY^Δ2^ was produced using guide RNAs flanking *Sry4A* and *Sry3C* (Figure 2A). This model harbors a deletion of most or all of ChrY distal to *Kdm5d*. Sequence analysis revealed a small inversion of the 3’ end of *Kdm5d* which was confirmed by PCR (data not shown) which can be used to distinguish the mutant ChrY^Δ2^ from wildtype ChrY using PCR (see Table 3). It should be noted that the WKY/Bbb assembly and OP905629 sequence assembled from the SHR/Akr strain differ in some repetitive regions from each other and from the reference BN (GRCr8) genome (data not shown), likely because of difficulty assembling the highly repetitive ChrY sequences or other inherent strain differences. Thus, it is currently not possible to determine a completely accurate map of the mutant ChrY^Δ^ in either of the SD XY^Δ^ models, although unique rearrangements were identified to distinguish the mutant ChrY^Δ^ from wildtype ChrY by PCR in each model.

### Production of transgenic lines

Integration of an *Sry* BAC clone harboring *Sry4A*, *Sry1*, and *Sry3C* (see Figure 3) into embryos of Sprague Dawley rats produced four lines of XYTG gonadal males that were fertile (Table 2) and transmitted the transgene to XX offspring (XXTG) with testes. Each of the four transgenic lines were backcrossed to Sprague Dawley Crl:CD(SD) for >10 generations to eliminate possible second transgenic integration sites or potential off-target anomalies in the genome. Each transgenic line produced frequencies of genotypes that were not different from expected Mendelian frequencies (Χ^2^ test, p > 0.05, Table 4A). The transgenic cross (Figure 1) produces three types of males: XXTG, XYWT, and XYTG.

**Figure 3.**
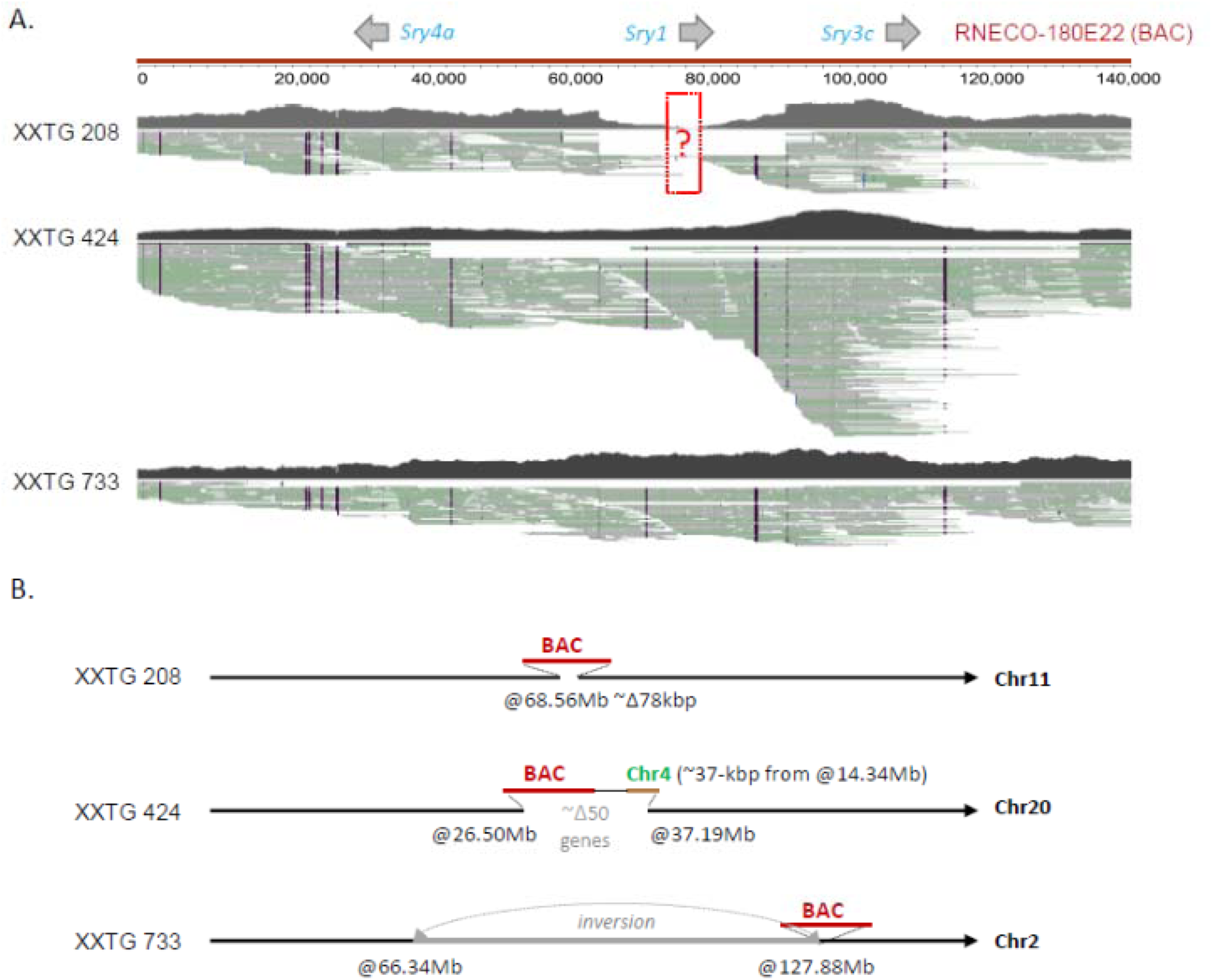
Genomic characterization of Sry BAC transgenic models by HiFi PacBio. A) Coverage and alignment of long-read data to RNECO-180E22. Line 208 may lack parts of the *Sry1* gene. B) Organization of the insertion sites of each transgene integration. The size of insertions, deletions, and inversions are not to scale.

A combination of paired-end Illumina sequencing and long-read PacBio HiFi sequencing was used to determine the genomic makeup of three lines (Figure 3). The fourth line, 737, was not studied because it was disqualified based on measurements of anogenital distance, see below. Comparison to the unannotated SR/JrHsd genome, which was derived from selective inbreeding of Sprague Dawley rats, allowed confirmation of the integration of the BAC DNA at unique autosomal locations in each line (Figure 3B). Line 208 harbors an integration on Chr11, which caused a ∼78-kbp deletion surrounding the integration site. No genes are annotated in this region in the GRCr8 BN rat reference genome (data not shown). Line 424 harbors a complex integration on Chr20 which also includes a small ∼37-kbp segment of Chr4. Integration of the transgene in this line resulted in a >10Mbp deletion on Chr20 involving dozens of genes. This loss of genes makes line 424 undesirable for use as part of an FCG analysis. Finally, line 733 harbors an integration of the BAC on Chr2 but caused a >60Mbp inversion in the middle of the chromosome. Sequence breakpoints were identified at various junctions in all three lines and confirmed using unique PCR primer sets (data not shown, see Table 3). FISH analysis of metaphase spreads confirmed the location of the BAC transgene on Chr11 (line 208) (Figure 4), and Chr2 (line 733) and Chr20 (line 424) (not shown)[38]. In each line, some *E.coli* DNA from the host bacteria used to propagate the BAC was detected (see Supplementary Figure 1). Based on KRAKEN2 assessment of broad species for liver RNAseq, the transgenic lines do not express *E. coli* transcripts above the background level found in wild-type rats. The *de novo* assembly of liver RNAseq reads primarily annotated mouse (102,365) and rat (55,046) spliced transcripts, with only three *E. coli* transcripts at below traditional cutoff values of 1 transcript per million (TPM), which occurred in wild-type and transgenic lines alike (Supplementary Figure 2).

**Figure 4.**
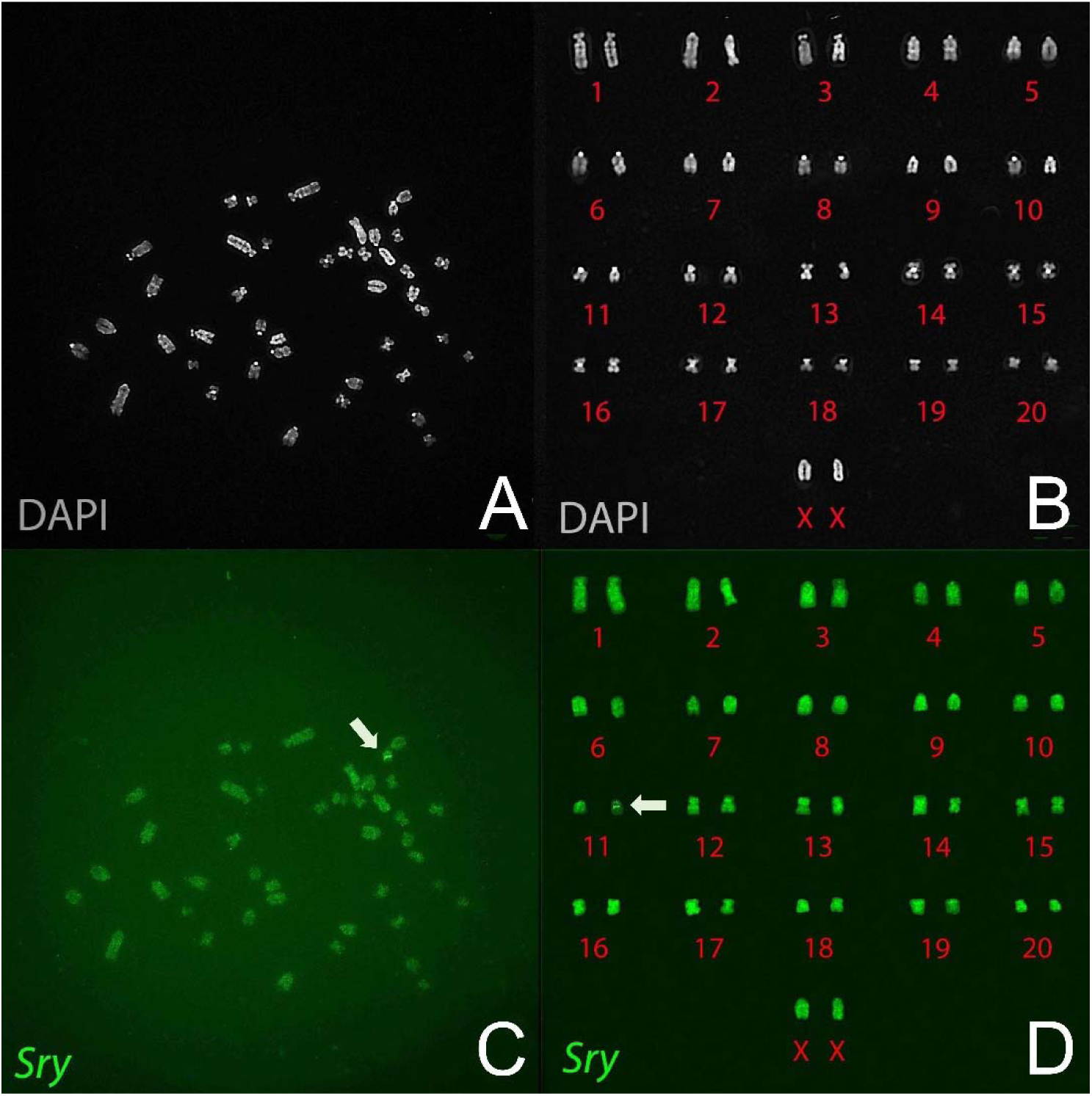
Localization of Sry-BAC transgene to Chr11 in metaphase chromosomes of an XXTG male of line 208. A, metaphase chromosomes labeled by DAPI. B, Sorted DAPI-stained chromosomes. C, the same chromosome spread labeled with the *Sry* BAC fluorescent probe. D, Sorted fluorescently labeled chromosomes. The arrows indicate presence of the *Sry* transgene on Chr11.

### Fertility of progeny

Three genotypes of atypical females (XO, XY^Δ^, XXY^Δ^), produced by these crosses, were mated to WT males. XO females produced litters with XO and XX females, and XY males (Table 2). XY^Δ^ females produced litters with XO, XX, XY^Δ^, and XXY^Δ^ females, and XY and XYY^Δ^ males. XXY^Δ^ females produced litters with XO (very rare), XX, and XXY^Δ^ females, and XY and XYY^Δ^ males (Figure 1, Table 2). When XY^Δ^ females were mated to XYWT males, their offspring were derived from X eggs 32-48% of time (i.e., sum of XX and XY offspring), Y^Δ^ eggs about 10% (XY^Δ^ offspring), XY^Δ^ eggs 39-50% (sum of XXY^Δ^ and XYY^Δ^), and “O” eggs 3-8% (XO offspring) (Table 2). These percentages show high rates of nondisjunction and suggest lower viability of eggs lacking an X chromosome [35]. XXY^Δ1^ females produced X and XY^Δ1^ eggs, the latter with frequencies in the range of 35-50% (Table 2). When the crosses just mentioned used an XYTG father, all of the genotypes are produced both without the transgene and as gonadal males possessing the *Sry* transgene (Table 2).

### Gonadal histology

External genitalia of transgenic males were comparable to WT males (Figure 5). XYWT and XYTG testes were comparable in size, but XXTG testes were smaller (Figure 5). Histological analysis showed that Sertoli cells in the XXTG testes formed seminiferous tubules, but the tubules lacked germ cells (Figure 6AB), as expected and shown previously in other species [54]. This is due to the presence of a second ChrX which is incompatible with spermatogenesis, coupled with the loss of Y-linked genes required for producing sperm [55, 56]. Lack of germ cells leads to a reduction in cord diameter [57] and in overall testis size as observed in other models of Sertoli-cell only syndrome [58]. All other gonadal cell types seen in the XYWT testes (such as Leydig cells, myoid cells and blood cells) were observed in approximately similar numbers in the XXTG testes. The testes of XYTG males were histologically similar to XYWT males (Figure 6AB).

**Figure 5.**
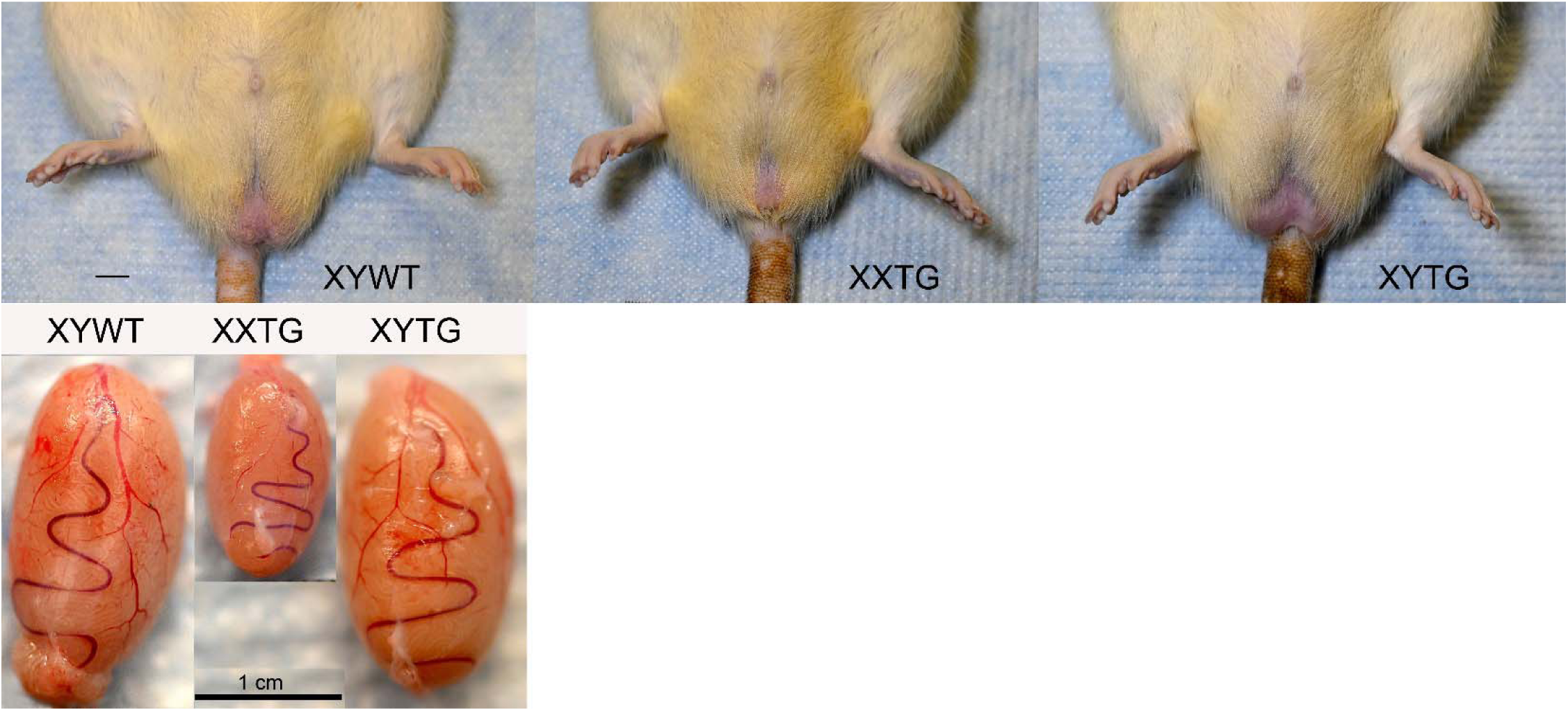
External genitalia and testis morphology. Above, External genitalia of XX and XY gonadal males are comparable. Below, testes of the three types of males. Testicular size was comparable in XYWT and XYTG males, but smaller in XXTG males. Photos from line 208 at postnatal day 60. Scale bars, 1 cm.

**Figure 6.**
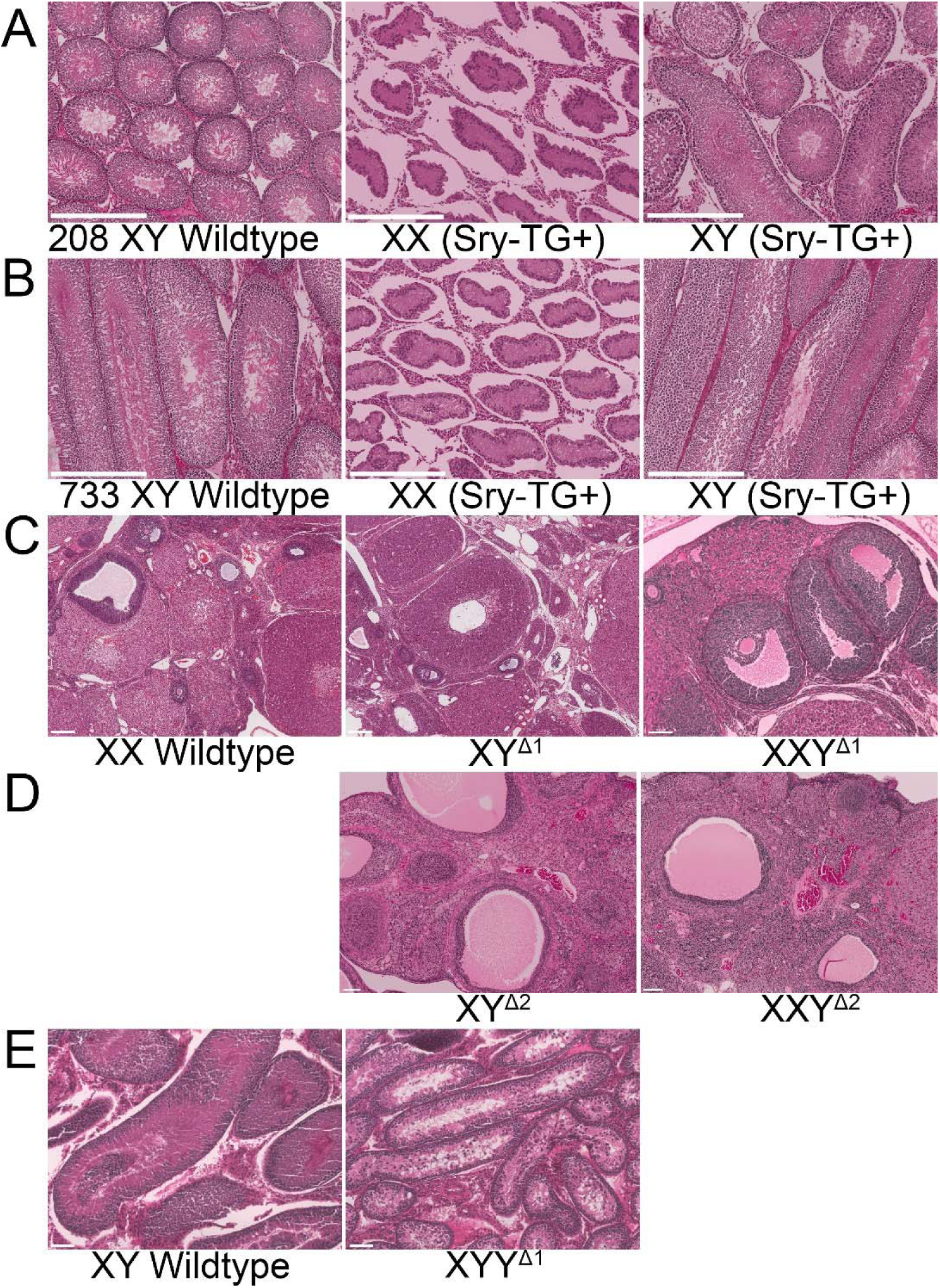
Gonadal histology. A, B. XY males of transgenic lines 208 and 733 (XYWT and XYTG) have comparable testicular morphology and distribution of cell types. XXTG males of both lines have seminiferous tubules and Leydig cells but lack cells of the germ cell lineage. C. Ovaries of XX, XY^Δ1^ and XXY^Δ1^ females have comparable histology, showing granulosa cells and all stages of folliculogenesis. D. Ovaries of XY^Δ2^ and XXY^Δ2^ show comparable histology, granulosa cells and all stages of folliculogenesis are present. E. XYWT and XXY^Δ1^ males show comparable testicular morphology with Sertoli cells forming seminiferous tubules and Leydig cells also present. XXY^Δ1^ males have spermatogonial stem cells, but lack spermatids and mature sperm. Hematoxylin and eosin staining. Scale bars are 100 μm.

Histological analysis of XXWT, XY^Δ1^, XXY^Δ1^ and XY^Δ2^ ovaries showed comparable morphology (Figure 6CD) and all stages of oocyte development present including primary and secondary follicles, antral follicles and corpora lutea. Likewise, development and organization of granulosa and theca cells were comparable in XXWT, XY^Δ1^, XXY^Δ1^ and XY^Δ2^ rats (Figure 6CD). Although the XY^Δ2^ females were significantly older than the XY^Δ1^ females at the time of histology collection, all stages of folliculogenesis were observed, suggesting that the genetic modifications are unlikely to cause premature ovarian failure or early infertility.

XYWT and XYY^Δ1^ testes were comparable in size. Histological analysis showed that Sertoli cells in the XYY^Δ1^ testes formed seminiferous tubules, and the tubules had spermatogonia and spermatocytes but lacked spermatids and mature sperm (Figure 6E). This is consistent with previous work in mice showing XYY males are infertile [59] due to chromosome asynapsis rather than the presence of the additional Y [60]. Other testicular cell types such as Leydig cells, myoid cells and blood cells were observed in XYY^Δ1^ testes (Figure 6E).

### Analysis of Sry expression in tissues

We measured expression of *Sry* transcripts using oligo-dT primers in a reverse transcription reaction, and qPCR with primers that amplify cDNAs from all *Sry* genes except *nonHMGSry*. Adult spleen, liver and gonad were assayed in males of transgenic lines 208 and 733, and from XXWT, XY^Δ1^, and XXY^Δ1^ females produced by XY^Δ1^ mothers (Figure 7). In all tissues, *Sry* expression varied by genotype (one-way ANOVA, p<0.00001, Table 4B). In spleen and liver, XYWT and XYTG males had higher levels of expression than XXTG males (TK, p < 0.05), as expected because the two former groups have a ChrY but XXTG males do not (Figure 7AB). *Sry* was expressed from the transgene in spleen and liver as shown in XXTG males (Figure 7AB). *Sry* expression might be expected to have been higher in XYTG than XYWT in these tissues because of the additional copies of *Sry* in the transgene, but that difference was not statistically significant (TK, p > 0.05). *Sry* was expressed in spleen and liver at comparable levels in XY^Δ1^ and XXY^Δ1^females (Figure 7), confirming the presence of functional *Sry* gene(s) on the Y^Δ1^ chromosome, and suggesting that *Sry* genes remaining on the Y^Δ1^ chromosome are normally expressed in those tissues in XYWT males.

**Figure 7.**
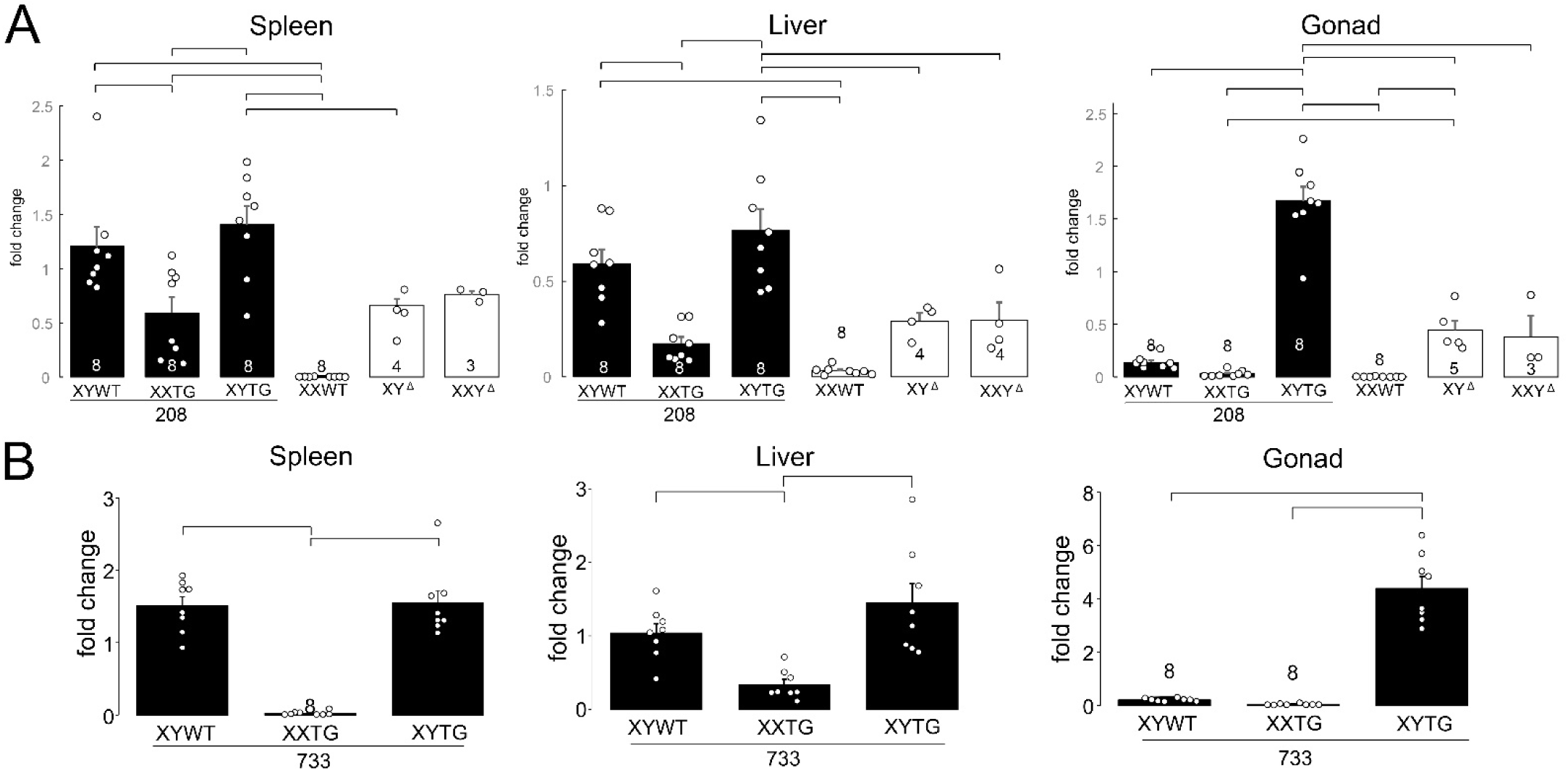
Expression of *Sry* mRNA in tissues measured by RTPCR. A. In transgenic line 208 and progeny of XY^Δ1^ females, expression varied significantly among the six genotypes (one-way ANOVA, p < 0.000001). Numerous groups differed in pairwise posthoc analyses (Tukey-Kramer, p < 0.05 or better) as shown by brackets. B. In transgenic line 733, three gonadal male groups showed significant variation (one-way ANOVA, p < 0.000001), and some groups differed from others (Tukey-Kramer p < 0.05) as shown by brackets. Black bars depict groups with testes, and white bars depict groups with ovaries.

In adult gonads, *Sry* was expressed in all genotypes except for XXWT females, but at much higher levels in XYTG than in other groups. The higher expression of *Sry* from XYTG testes, relative to XYWT testes, might imply that expression from the transgene was much higher than from the endogenous ChrY. However, expression from the transgene in XXTG males was quite low, undermining that simple conclusion. Thus, factors on ChrY and in the transgene might interact to regulate *Sry* expression in XYTG but not XXTG groups. Expression of *Sry* in ovaries of XY^Δ1^ and XXY^Δ1^ rats again attests to the presence of *Sry* on the Y^Δ1^ chromosome. The expression of *Sry* in ovaries is unprecedented and perhaps paradoxical, because ovaries normally lack *Sry*. Thus, any cell type, even ovarian cells, may possess mechanisms that activate expression of *Sry* or other ChrY genes. Our analysis indicates that *Sry2* is present on Y^Δ1^ (Figure 2). *Sry2* appears to be widely expressed in many tissues of WT males and is unlikely to be testis-determining [61], and may account for some of the *Sry* expression seen in Figure 7.

### Growth curves

We compared the body weight increase of the four transgenic lines from 3 to 8 weeks of age (Figure 8). In general, the XXTG males grew within the male range, but at slightly slower rates than XY groups. These differences were significant only in some lines. A repeated measures two-way ANOVA with factors of genotype (3 levels) and age (6 levels) showed a main effect of age (p<0.00001, Table 4C) in all lines, and either no effect of genotype (lines 733 or 424), or a main effect of genotype (line 737, p=0.05, Table 4C), or a genotype by age interaction (line 208, p<0.0008, Table 4C). Because the two XY genotypes (XYWT and XYTG) were not statistically different, we combined them and used a two-way ANOVA (factors of genotype (2 levels) and age (6 levels) to test for XX vs. XY differences. In line 737, the main effect of genotype was significant (p<0.008, Table 4C). The XX-XY difference showed a significant interaction with age in line 208 (p<0.0003. Table 4C), line 733 (p<0.05, Table 4C), and line 737 (p<0.003, Table 4C). In line 424, the XX-XY difference was not significant. At week 8, a comparison of the combined XY groups with the XXTG group revealed a significant difference only for line 208 (p=0.034, Table 4C).

**Figure 8.**
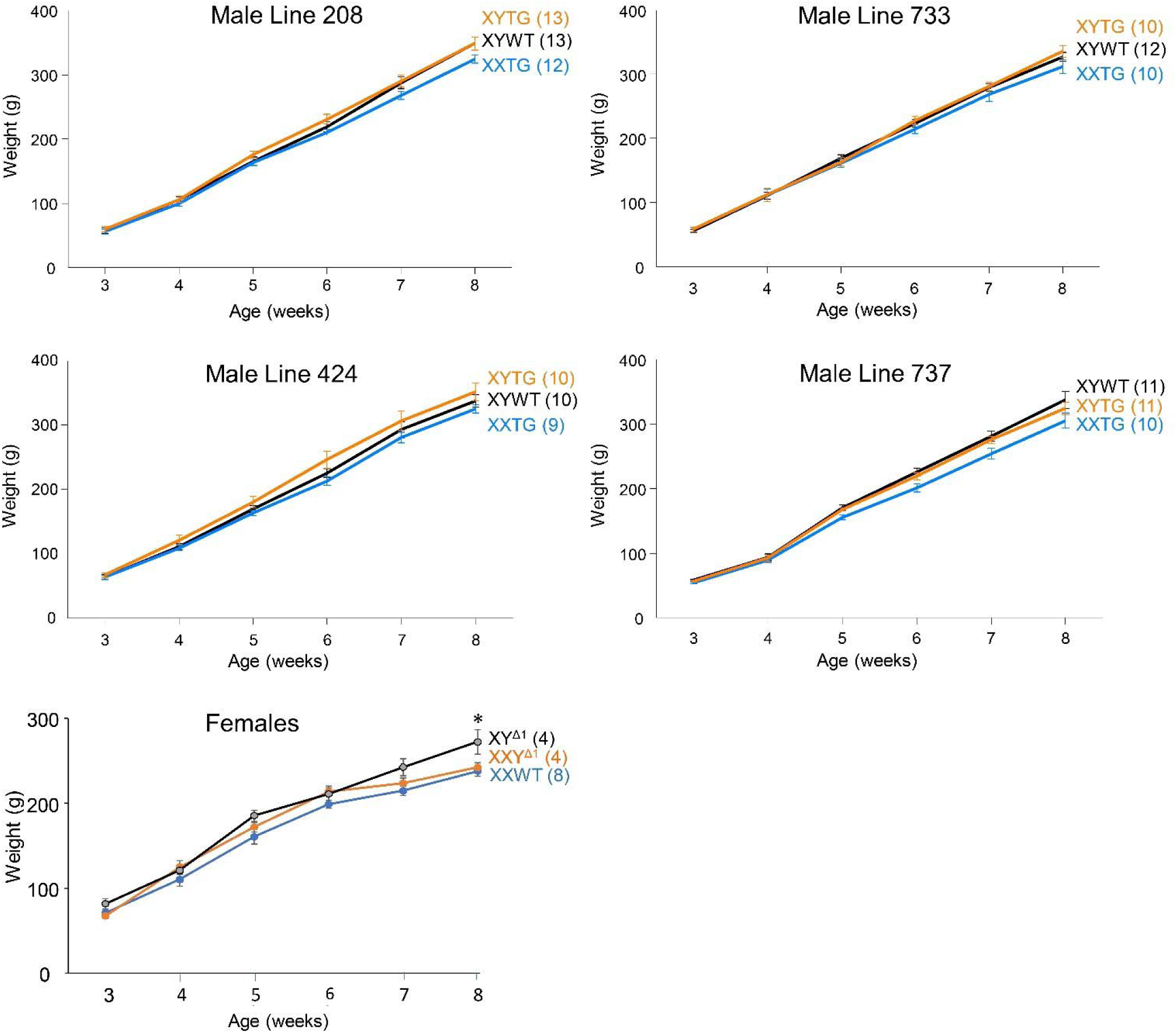
Growth curves. Graphs show growth in genotypes from 3 to 8 weeks of age in backcross generation 4 (females) or 7 (males). XYTG males grew at rates that were not statistically different from those of XYWT males, but XXTG males grew at slightly lower rates. The two XY groups were combined for statistical comparison to XXTG males, using a two-way ANOVA with factors of sex-chromosome complement (XX vs. XY) and age (repeated measure). Differences in body weight of XX and XY males were shown by a main effect of sex-chromosome complement (line 737, p=0.008), or because of a significant interaction of sex chromosomes and age (line 208, p<0.0003); line 733, p<0.05; line 737, p<0.003). In line 424, no significant effect of sex chromosomes was detected. In females, a two-way ANOVA with factors of sex-chromosome complement (XX vs. XY) and age (repeated measure) showed a significant interaction of sex-chromosome complement and age (p<0.05). At 8 weeks, the female groups differed in body weight (one-way ANOVA, p<0.04) because of greater weight of XY^Δ1^ females than XX females (Tukey-Kramer, p<0.05). Group size is indicated in parentheses.

Body weight among three female groups (XXWT, XY^Δ1^, XXY^Δ1^) showed a significant interaction with age over 3-8 weeks (p<0.05, two-way repeated measures ANOVA, Table 4C). At 8 weeks of age weight varied significantly among the groups (p<0.04, one-way ANOVA, Table 4C) because XY^Δ1^ females weighed more than XX females (TK, p<0.05).

### Organ weights

In backcross generation 7, we measured weights of kidney, spleen, and testes of male genotypes in all lines (Figure 9). Testis size differed among genotypes (one-way ANOVA, p<0.000001, Table 4D), because testes were smaller in each line in XXTG relative to XYWT and XYTG (TK, p<0.001). There were no significant differences in kidney and spleen weights among the three male groups in any line (one-way ANOVA, p>0.05, Table 4D). Representative data are shown for line 208 in Figure 9.

**Figure 9.**
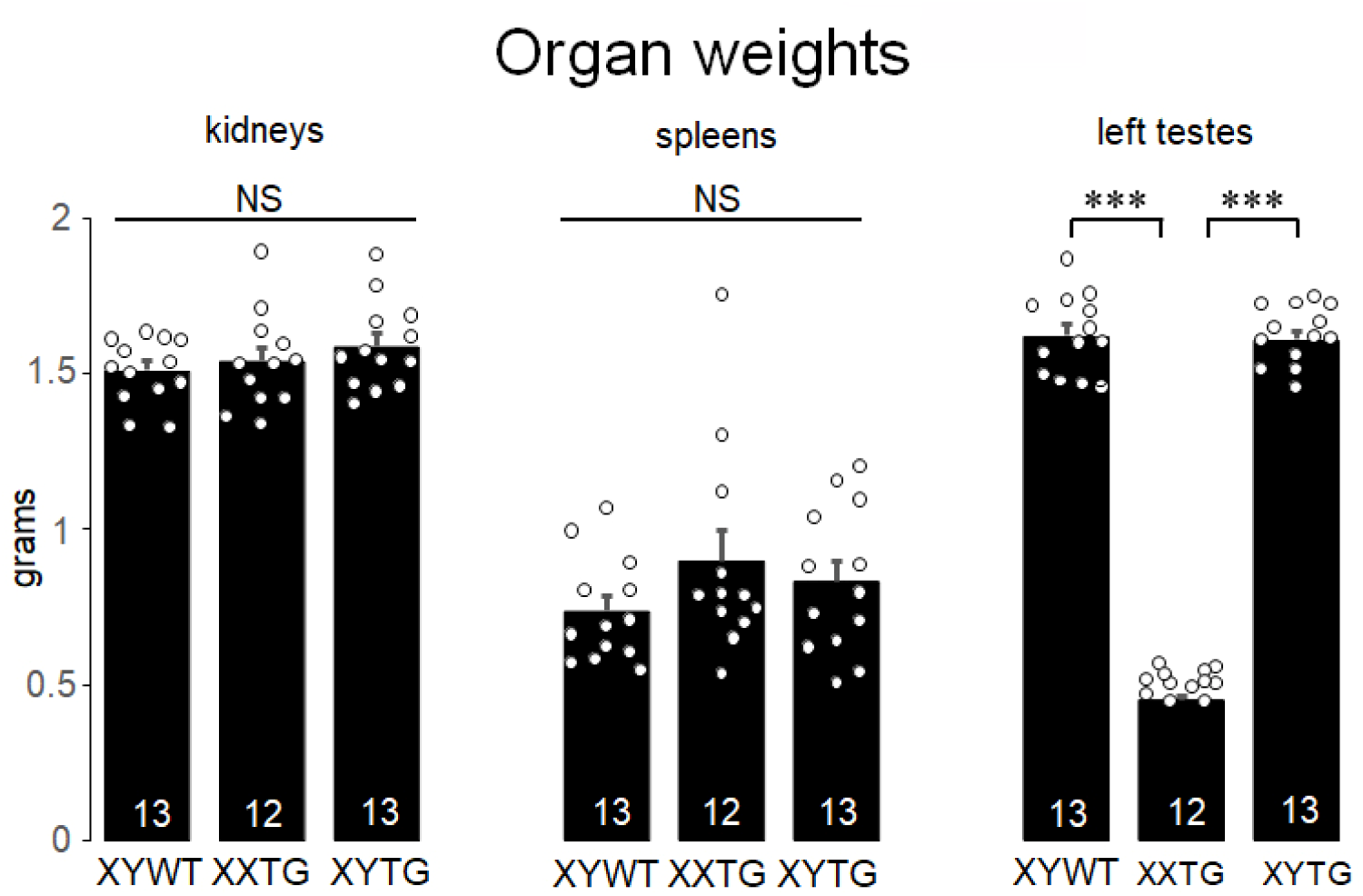
Organ weights. The weight of kidneys, spleens, and testes from line 208 at postnatal day 60, in backcross generation 7. There was no significant effect of genotype on kidney or spleen weight. Testis weight varied by genotype (one-way ANOVA, p<0.000001) because XXTG testes weighed significantly less than those of either XY group (*** Tukey Kramer, p<0.001). In all figures, histograms indicate mean, error bars are S.E.M., and group size is shown in each bar. NS, not significant.

### Anogenital distance

The anogenital distance, which is sensitive to prenatal levels of gonadal androgens, varied by genotype (one-way ANOVA, p<0.00001, Figure 10, Table 4E) reflecting the difference in AGD between rats with testes vs. ovaries. All male groups differed from all female groups (TK, p<0.001) but not from each other (TK, p>0.05). In line 737, the AGD of XXTG males was somewhat smaller than those of XYWT (TK p<0.01). In litters from XXY^Δ1^ mothers and XYTG fathers, the effect of genotype was also significant (one-way ANOVA, p<0.00001, Table 4E), but no significant differences were found among male groups or between the female groups (one-way ANOVAs, *p*s>0.25). The comparable AGD among same-gonad groups suggests that in all lines except 737, prenatal androgen levels are comparable among groups with testes, and among groups with ovaries.

**Figure 10.**
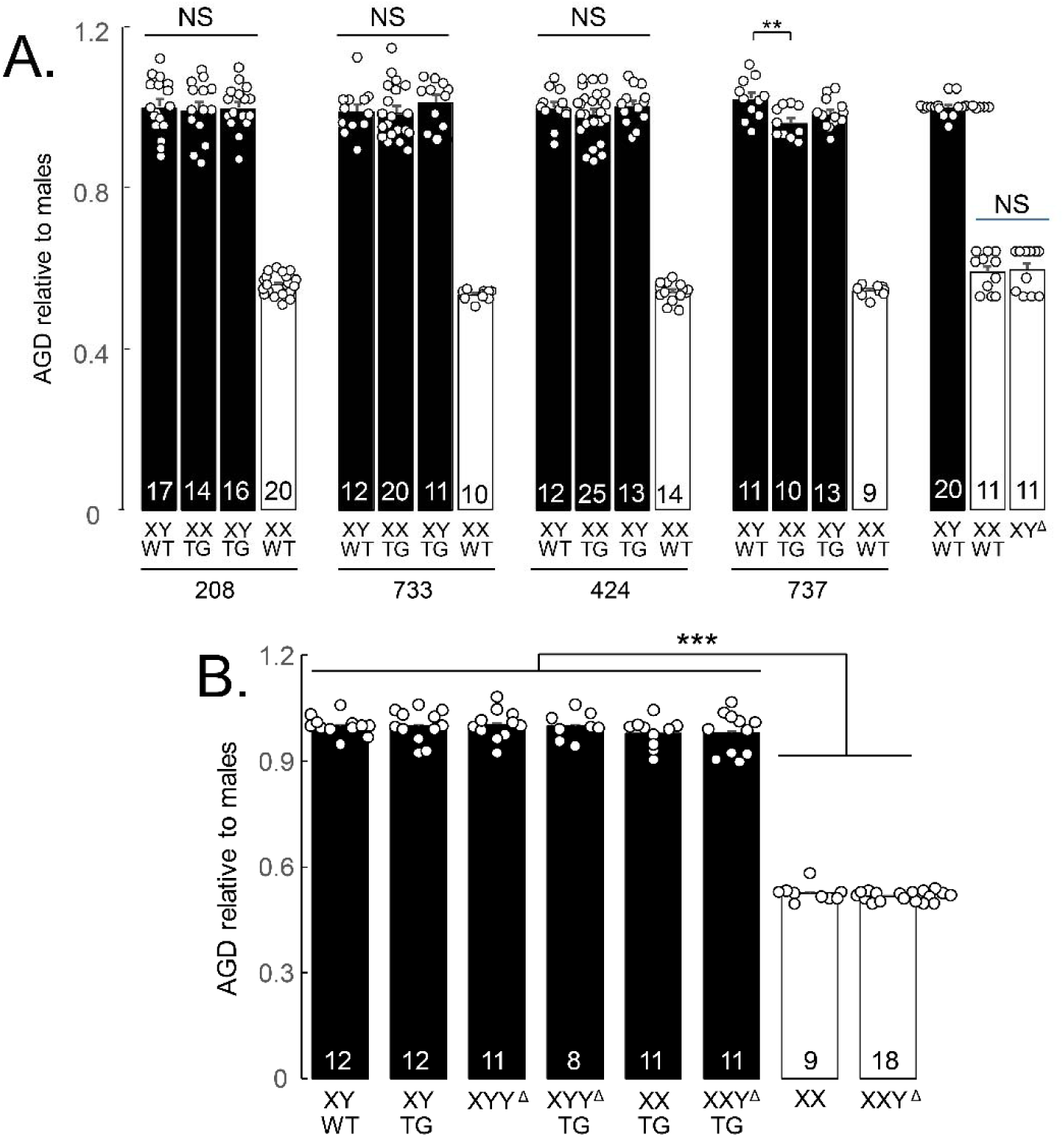
Anogenital distance. A. AGD was measured at P6-7 and expressed relative to mean AGD for XYWT males in the litter. In all lines, AGD differed among groups (one-way ANOVA, p<0.00001) because of a consistent difference in AGD between rats with testes or ovaries (Tukey-Kramer, p<0.001). In lines 208, 733, and 424, male genotypes did not differ in AGD. In line 737, XXTG had lower AGD than XYWT (Tukey-Kramer, **p<0.01). XX and XY gonadal females did not differ significantly in AGD. NS, not significant. B. AGD was measured in the same way in litters of the cross XXY^Δ1^ x XYTG. Groups differed in AGD (one-way ANOVA, p<0.00001). All groups with testes had greater AGD that all groups with ovaries (TK, p<0.05), but all differences between other group dyads were not significant.

### Hormone levels

In backcross generation 4, we measured serum testosterone, LH, and FSH levels at ∼P60, in lines 208 and 424 (Figure 11). Serum testosterone levels did not differ significantly among male groups in each line (one-way ANOVA, p>0.05, Table 4F). LH levels varied significantly with genotype in line 208 (one-way ANOVA, *p*<0.007, Table 4F) because XXWT levels were significantly lower than in XYWT and XXTG males (TK, *p*<0.05). LH levels did not vary significantly in genotypes from line 424 or in progeny of XY^Δ1^ females (one-way ANOVA, p>0.14, Table 4F).

**Figure 11.**
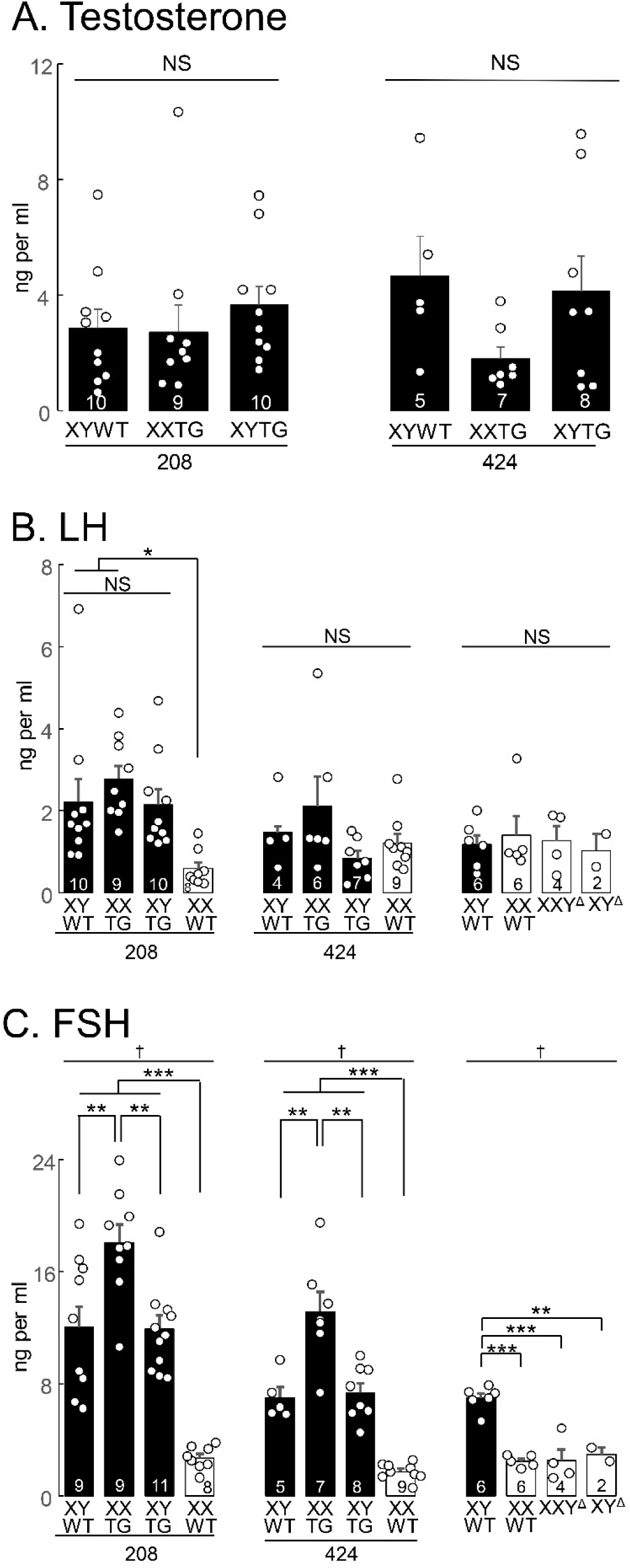
Levels of hormones. A., serum testosterone did not differ significantly among male groups in lines 208 or 424 (one-way ANOVA). B. LH levels did not differ significantly among groups in line 424, or among progeny of XY^Δ^ females. LH levels varied significantly with genotype in line 208 (one-way ANOVA, p<0.007) because XXWT levels were significantly lower than in XYWT and XXTG males (Tukey-Kramer, * p<0.05). C. FSH levels in lines 208 and 424 differed across genotypes (one-way ANOVA, † p<0.00002). XXWT females had lower levels than each of the male groups (***TK for 208 p<0.001), and the XXTG group had higher levels than XYWT or XYTG groups (Tukey-Kramer, ** p<0.01). FSH levels in litters of XY^Δ^ females (right side) were significantly different among groups (one-way ANOVA † p<0.00002), because each female group differed significantly from the XYWT group (TK, *** p<0.001) but not from each other. Measurements were made in backcross generation 4 at P60. NS, not significant.

FSH levels in lines 208 and 424 differed across genotypes (one-way ANOVA, p<0.00002). XXWT females had lower levels than each of the male groups (line 208, TK p<0.001), and the XXTG group had higher levels than XYWT or XYTG groups (TK, p<0.01). FSH levels in litters of XY^Δ1^ females were significantly different among groups (one-way ANOVA, p<0.00002), because each female group differed significantly from the XYWT group (TK, p<0.001) but not from each other.

We measured serum testosterone levels in males of all transgenic lines at backcross generation 7 (Figure 12), but found no significant differences among male groups in each line (one-way ANOVA, Table 4F).

**Figure 12.**
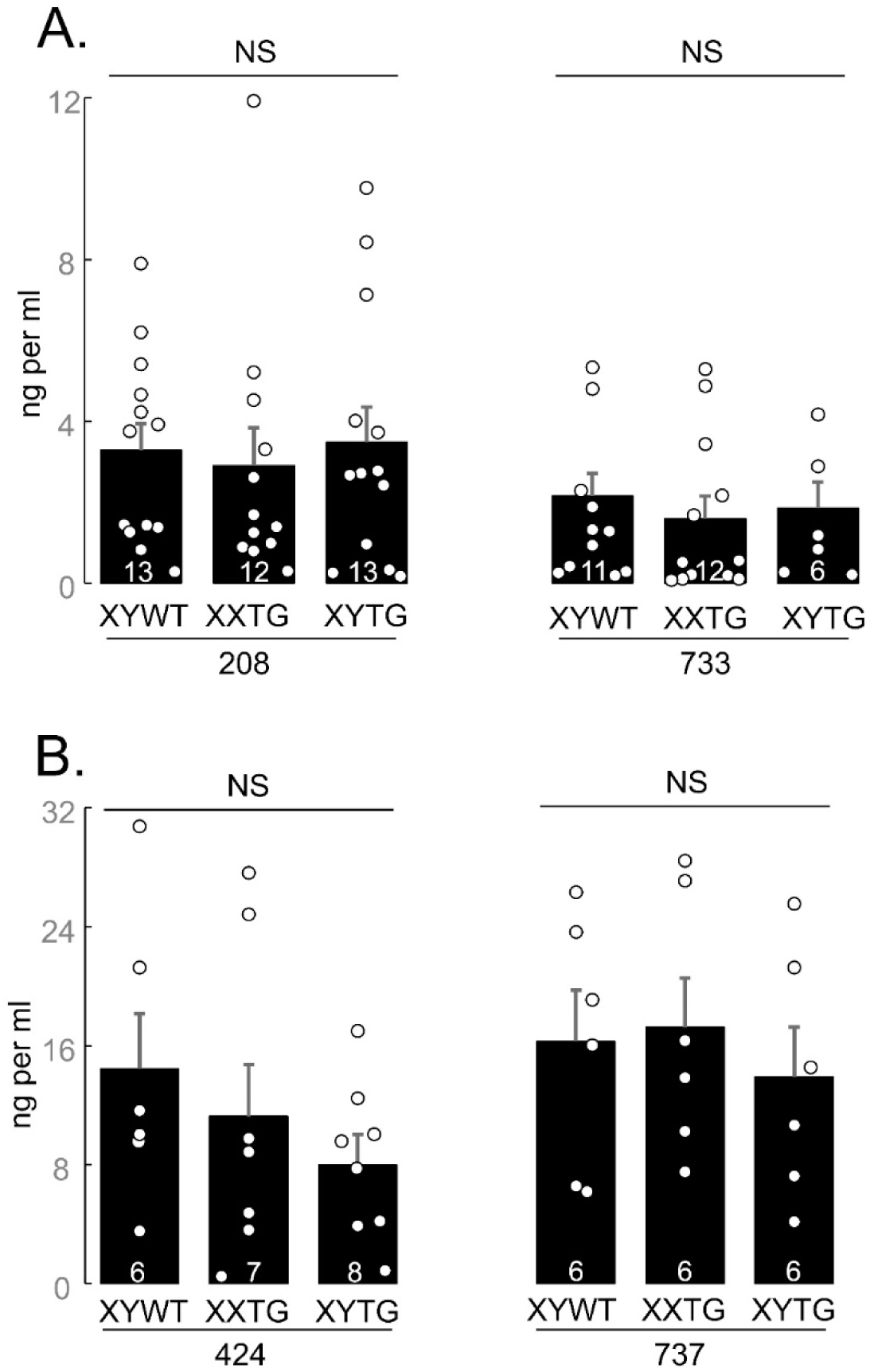
Serum testosterone in backcross generation 7 at P60. No significant differences (NS) were found among genotypes in all transgenic lines (one-way ANOVA).

### Analysis of central nervous system (CNS) sexual dimorphisms: SNB, DLN, SDN-POA, and AVPV

Analysis of several brain attributes known to exhibit sexual dimorphism all showed differences between gonadal males and gonadal females. Dependent variables included the number and size of SNB and DLN motoneurons (Figures 13, 14), volume, cell number and cell size of SDN-POA CALB-ir neurons (Figure 15), and number and size of TH-ir cells in the AVPV (Figure 16). With minor variation, the three groups of gonadal males had comparable values for these dependent variables, which were significantly different than the values for the three groups of gonadal females, which also did not differ from each other. A one-way ANOVA revealed a significant effect of genotype in each case (p <0.000001, except p <0.02 for DLN soma size, Table 4G).

**Figure 13.**
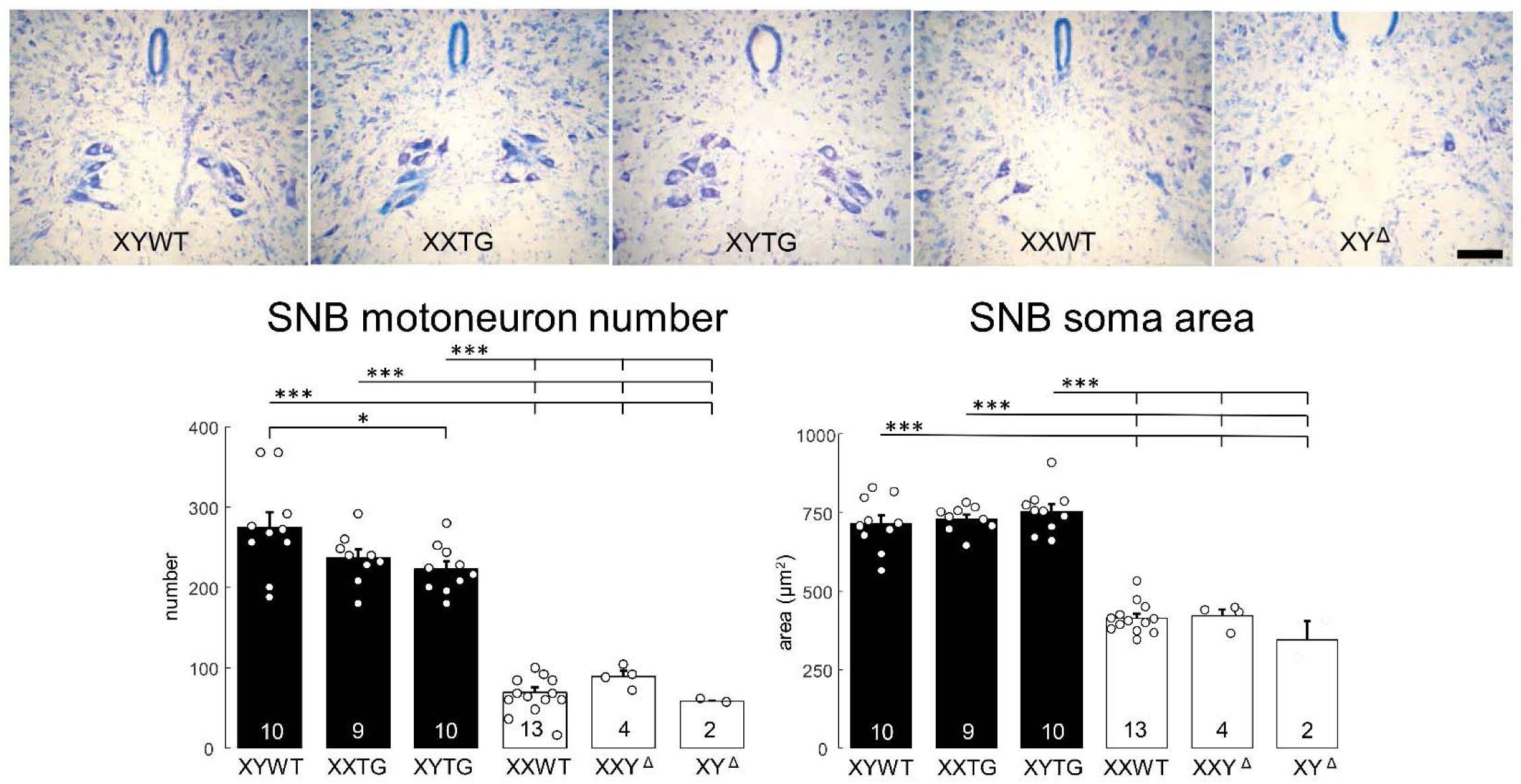
SNB morphology. Above, digital micrographs of thionin-stained transverse sections through the lumbar spinal cord of line 208 and progeny of XY^Δ1^ females, showing the medially located Spinal Nucleus of the Bulbocavernosus (SNB), with more neurons in groups with testes than in those with ovaries. Scale bar = 100 µm. Below, counts and soma areas of SNB motoneurons. Both motoneuron number and soma size varied significantly by genotype (p<0.000001), and all gonadal male groups were significantly different from all gonadal female groups (Tukey-Kramer, *** p<0.001). SNB number was lower in XYTG than XYWT (Tukey-Kramer, p<0.05). There was no significant variation between groups differing in sex-chromosome complement.

**Figure 14.**
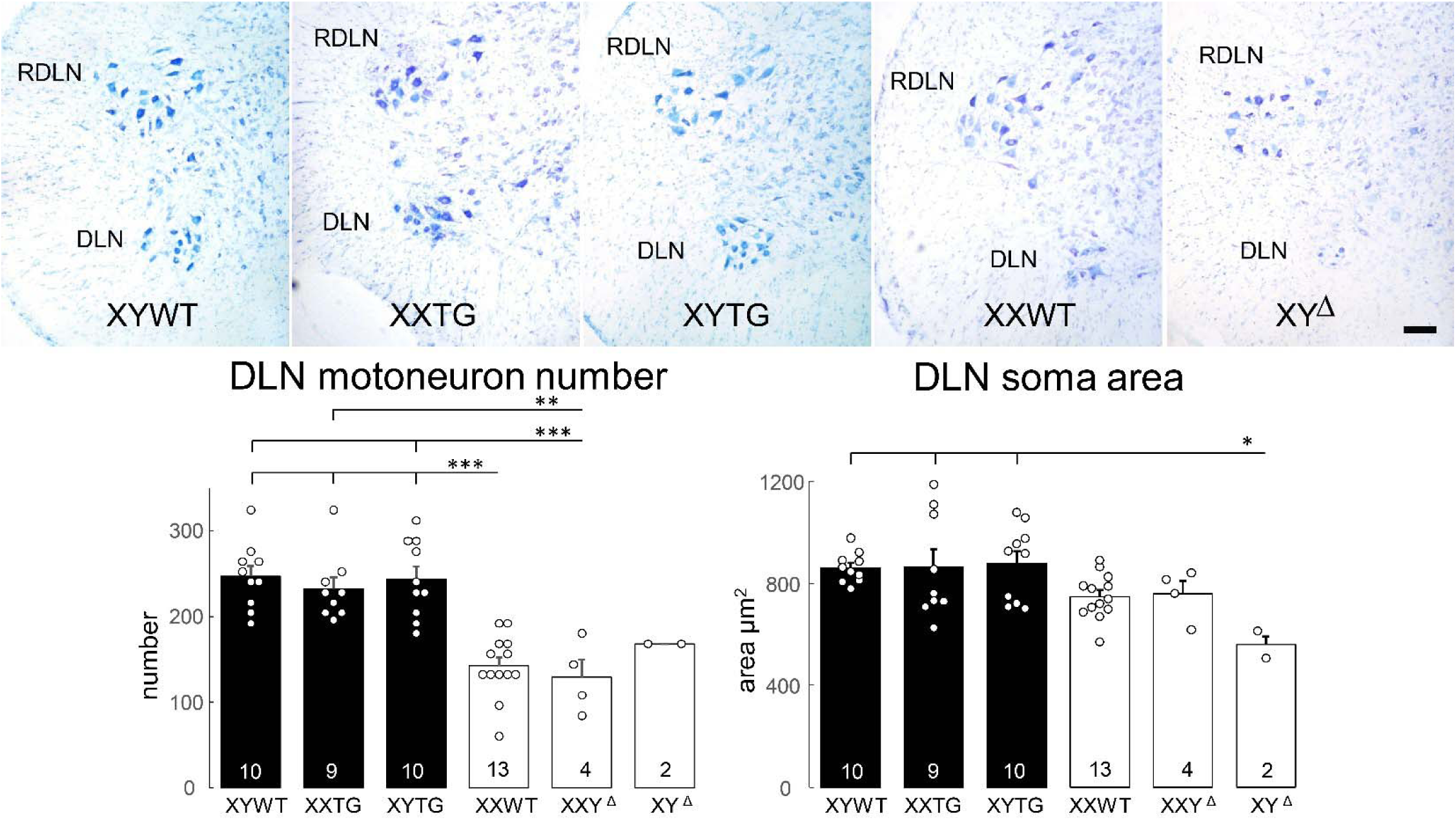
DLN morphology. Above, digital micrographs of thionin-stained transverse sections through the lumbar spinal cord show the laterally located Dorsolateral Nucleus (DLN) in line 208 and progeny of XY^Δ1^ females, showing more motoneurons in groups with testes than with ovaries. Also shown is the Retrodorsal Lateral Nucleus (RDLN), containing motoneurons that innervate an intrinsic muscle of the foot. Scale bar = 100 µm. Below, counts and soma areas of DLN motoneurons. The variation among groups was statistically significant (one-way ANOVA, p<0.000001 for DLN number, p< 0.02 for DLN area). For DLN number, all gonadal male groups differed from XXWT females and XXY^Δ1^ females (TK, p<0.01). For DLN soma area, XY^Δ1^ females differed from all gonadal male groups (TK<0.05). For DLN soma area, a two-way ANOVA with factors of gonad type (ovaries vs. testes) and sex-chromosome complement (XX vs. XY) revealed a main effect of gonad type (p< 0.001) but no significant effect of sex-chromosome complement or interaction (p>0.05).

**Figure 15.**
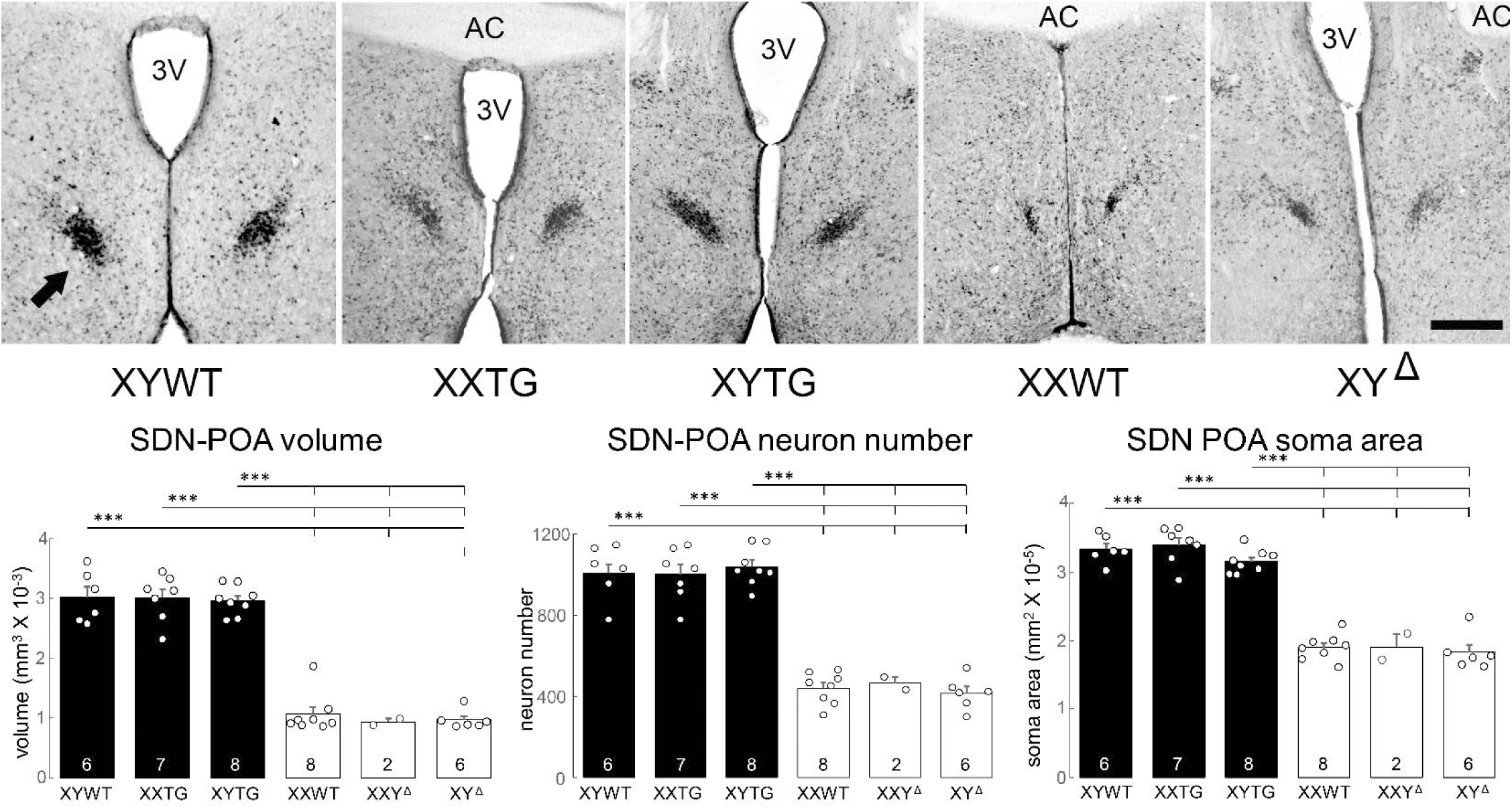
SDN-POA morphology. Above, digital micrographs of calbindin-immunoreactive cells in the Sexually Dimorphic Nucleus of the Preoptic Area of the hypothalamus (arrow) with nearby anterior commissure (AC) and third ventricle (3V) in line 208 and progeny of XY^Δ1^ females. The SDN-POA is larger in groups with testes than in those with ovaries. Scale bar = 300 μm. Below, measurements of SDN-POA volume, neuron number, and immunoreactive cell area. For each type of measurement, variation among groups was statistically significant (one-way ANOVA, p < 0.000001), and each group with testes was significantly different from each group with ovaries (Tukey-Kramer, *** p < 0.001). There was no significant variation between groups differing in sex-chromosome complement.

**Figure 16.**
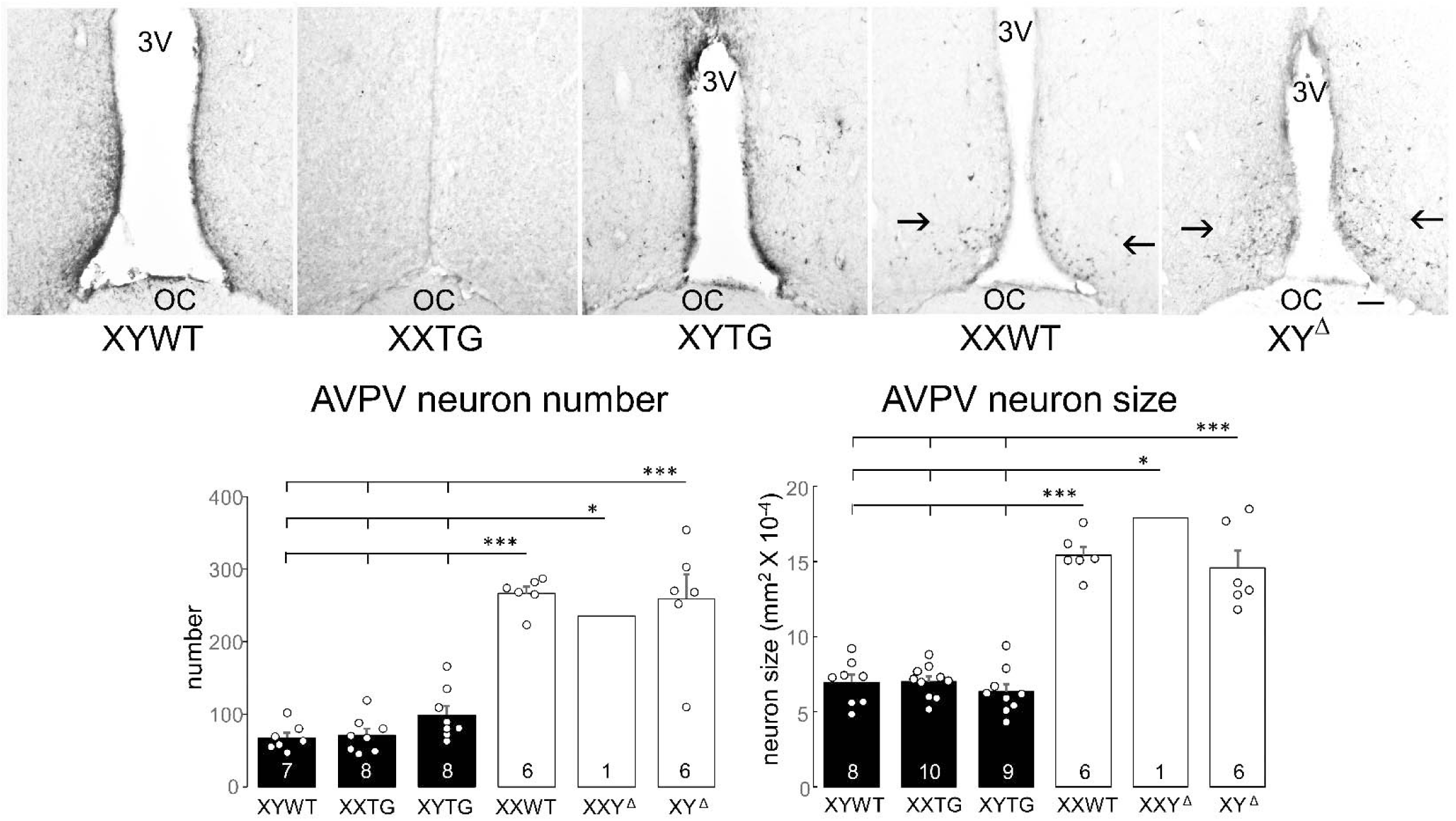
AVPV morphology. Above, digital micrographs of tyrosine-hydroxylase-immunoreactive neurons in the hypothalamic Anteroventral Periventricular Nucleus (AVPV) (arrows) adjacent to the third ventricle (3V) and above the optic chiasm (OC) in line 208 and progeny of XY^Δ1^ females. Scale bar =100 μm. Many more TH-ir neurons are found in rats with ovaries than in rats with testes, independent of sex-chromosome complement. Below, measures of numbers and size of TH-ir neurons. In both variables, variation among groups was significantly different (one-way ANOVA, p < 0.000001). Each gonadal female group had more neurons than each gonadal female group (Tukey-Kramer, *** p < 0.001 of *p<0.05). There were no significant differences between groups differing in sex-chromosome complement.

Differences between all pairs of gonadal male and female groups were significant (TK, p<0.05), except that for DLN soma area only the XY^Δ1^ female group differed from the three male groups (TK, p<0.05). In that case, however, a two-way ANOVA with factors of sex-chromosome complement (XX vs. XY) and gonad type (ovaries vs. testes) indicated that DLN soma size was significantly different between gonadal males and females (p<0.001, Table 4G) but not between XX and XY groups. In these CNS variables, no differences were found when comparing XX vs. XY groups with the same type of gonads (TK, p>0.05). These results support the conclusion that these sexual dimorphisms in rat CNS are caused by different secretions of gonadal hormones, confirming conclusions of previous reports based on manipulations of levels of gonadal hormones in WT males and females [48, 49, 62–65]. The current analysis also found no evidence that sex-chromosome complement influences these sexual dimorphisms. However, XYWT males had a larger number of SNB motoneurons than XYTG males (TK p<0.05), raising the possibility that the transgene produces some additional effect in XY males.

## Discussion

We sought to develop broadly useful rat models to distinguish sex-chromosomal and gonadal hormonal effects that contribute to sex differences in physiology and disease. We integrated a functional BAC *Sry* transgene into an autosome in four lines, resulting in XYTG male rats that produce XXTG offspring with testes. We also used CRISPR technology to disrupt ChrY regions required for testis determination, producing XY^Δ^ gonadal females that are fertile. The transgenic breeding lines produced XXTG gonadal males that can be compared to littermate XY males (XYWT and/or XYTG), to measure the differential effects of XX vs. XY sex chromosomes in a testicular hormonal environment (Figure 1, Tables 1 and 2). Breeding the XY^Δ^ females to XYWT males produced XXWT and XY^Δ^ females that can be compared to measure the differential effects of XX and XY sex chromosomes in an ovarian hormonal environment. The two comparisons of XX and XY rats, with the same type of gonad, offer a unique opportunity to determine whether sex differences in any rat phenotype are the result of XX vs. XY sex-chromosome complement, not mediated by gonadal secretions. The genotypes produced also allow the comparison of rats with different gonads, but with comparable sex chromosomes (either XX or XY), to determine whether sex differences in rat phenotypes are caused by gonadal hormones.

These comparisons are similar (but not identical, see below) to those afforded by the Four Core Genotypes (FCG) mouse model, which has been used extensively to uncover sex-chromosome effects that contribute to sex differences in many mouse phenotypes [3, 4, 7]. Because rats offer advantages for study of a different set of phenotypes than mice, the advent of an FCG-like rat model opens the door for expanded research on sex-chromosome effects. The ability to study sex-chromosome effects relatively easily, in a second species, will facilitate the process of evaluating species differences and evolution of sex-chromosome effects.

### Utility of sex-chromosome aneuploid rats

The nondisjunction of ChrX and ChrY during oogenesis in XY^Δ^ females leads to the production of “O”, X, Y, and XY^Δ^ eggs, and therefore three groups of sex-chromosome aneuploid offspring when the father is XYWT: XO females, XXY^Δ^ females, and XYY^Δ.^males (Tables 1 and 2, Figure 1). Other sex-chromosome aneuploid groups can be produced when XY^Δ^ or XXY^Δ^ females are mated with XYTG males: XOTG and XXY^Δ^TG and XYY^Δ^TG transgenic males (Tables 1 and 2, Figure 1). These groups offer important comparisons of rats differing in the number of ChrX or ChrY. If a sex-chromosome effect is discovered when comparing XX and XY rats with the same type of gonad, further tests can reveal whether the sex-chromosome effect is caused by the number of ChrX or ChrY. Comparing rats with different numbers of ChrX (XO vs. XX females or males, XY vs. XXY males) allows detection of ChrX effects. Comparing rats with different numbers of ChrY (XO vs. XY males, XX vs XXY females, XY vs XYY males) tests whether a sex-chromosome effect is caused by ChrY dose. These comparisons are similar to those possible using the XY* mouse model, which has been used as part of a fruitful strategy ultimately to identify specific X or Y genes that contribute to sex differences in mouse phenotypes [4, 7, 66–68]. The production of sex-chromosome aneuploid rats also has similarities to the Sex Chromosome Trisomy mouse model [69, 70].

### Functions of *Sry* genes

The current results contribute to an understanding of the function of some of the 11 different *Sry* genes in laboratory rats, whose diverse functions are unknown [32, 34]. The testis-determining effect of the BAC transgene indicates that *Sry4A, Sry1*, and/or *Sry3C* are sufficient, individually or in combination, to cause testis development in an XX rat. Several *Sry* genes remain on the Y^Δ^ chromosomes, based on analysis of the chromosome sequence and expression of *Sry* in tissues from XY^Δ^ females (Table 2, Figure 7). These remaining *Sry* genes are not sufficient by themselves, and in the numbers of copies remaining on the Y^Δ^ chromosomes, to cause testis development. However, these *Sry* genes may nevertheless contribute to testis differentiation in XYWT males but require the presence of the *Sry* genes deleted from the Y^Δ^ chromosomes. If the various *Sry* genes have similar functions, the formation of testes in XXTG but not XY^Δ^ groups could simply reflect a different timing or greater level of expression of *Sry* in the embryonic XXTG gonadal ridge, compared to XY^Δ^, at the onset of testis development. Alternatively, the different encoded genes could have different functions.

*Sry* was expressed in adult spleen and liver in virtually all genotypes that possess either the *Sry* transgene (encoding only *Sry4A*, *Sry1*, and *Sry3C*), or ChrY, or the Y^Δ1^ chromosome (which lacks multiple *Sry* genes including *Sry4A*, *Sry1*, and *Sry3C*) (Figure 7). Thus, these tissues likely typically express multiple *Sry* genes in XYWT males. The pattern confirms previous reports of extensive expression of *Sry* in non-gonadal tissues of rats [61]. Although *Sry2* may be an *Sry* gene expressed widely or ubiquitously [61], the present results imply that it is not the only *Sry* gene expressed in adult tissues. Because different genotypes expressed varying levels of *Sry* in diverse tissues, and expressed different *Sry* genes (Figure 7), these variations in *Sry* expression could contribute to group differences in emergent phenotypes tested using the rat models.

### Analysis of hormones

Sex differences are caused by two major classes of proximate sex-biasing factors, gonadal hormones and sex chromosomes [1]. Our intent was to create rat models that separate the effects of these two factors. Separating the two types of effects works best if same-gonad XX and XY rats have similar levels of gonadal hormones, so that group differences in hormone levels do not confound differences caused by sex-chromosome complement. We used several approaches to compare levels and effects of gonadal hormones in the rat models. (1) We found that AGD was similar in XX and XY gonadal males for three of four transgenic lines (208, 733, and 424), and similar in XX and XY^Δ^ and XXY^Δ^ gonadal females (Figure 10). This result implies that prenatal levels of androgens, which cause the sex difference in AGD [71–73], are comparable in these lines. Thus, the long-lasting prenatal “organizational” effects of testicular androgens, which cause numerous sex differences in non-gonadal phenotypes [74], are probably comparable in XX and XY gonadal males. The exception was line 737, in which AGD was well within the male range in XXTG males but slightly lower than that of XY males. That discovery led to termination of further work with line 737. (2) Measurement of circulating levels of testosterone in adult males showed that XXTG males had comparable levels of plasma testosterone, relative to XY males, in all four lines measured at about 60 days of age (Figures 11 and 12). (3) We examined several classic CNS sexual dimorphisms in line 208, including the SNB, DLN, SDN-POA, and AVPV (Figures 13-16), which are considered sensitive bioassays for levels of testicular hormones, because various sexually dimorphic attributes of these neural systems (cell number and size, nucleus volume) are caused by sex differences in prenatal and/or postnatal levels of testicular androgens [48, 49, 62–65]. All of these CNS regions in line 208 show a common pattern in measures of cell number, cell size, or nucleus volume. XXTG males are similar to XYWT males and strikingly more masculine than the three groups of females (XXWT, XY^Δ^, and XXY^Δ^), which are similar to each other (Figures 13-16). This pattern suggests that the levels of hormones that cause sexual differentiation of these tissues are comparable among gonadal male groups, and also in the female groups. The one footnote to this conclusion is that for cell number in the SNB, XYTG males had slightly fewer SNB motoneurons than XYWT males, raising the possibility of a small effect of the transgene on hormone levels in XY males.

### Body weight

In rats, the large sex difference in body weight (male > female) is controlled by perinatal and adult effects of ovarian and testicular hormones that influence feeding and have effects on adipose and other tissues regulating metabolism [75–77]. For example, gonadectomy at birth eliminates sex differences in rat body weight [77]. In rats studied here, animals with testes had body weights in the range typical of XYWT males, and animals with ovaries had body weights in the range of XXWT females, confirming the long-standing conclusion that sexual differentiation of body weight is controlled by gonadal hormones in rats (Figure 8). However, we found evidence in both gonadal males and females for small sex-chromosome effects (XY>XX) at least at some ages. Further work is needed to evaluate this possibility. The direction and size of the sex chromosome effect on body weight in rats are different from those in FCG and XY* mice that we previously reported [8, 18].

### Gonadotrophin levels

We detected gonadal hormone effects on LH and FSH levels, and sex-chromosome effects on FSH levels (Figure 11). LH levels were higher in gonadal males than females in line 208, a result that was not confirmed in line 424 or in progeny of XY^Δ^ females. FSH levels were robustly higher in XXTG males than in XYWT or XYTG males in lines 208 and 424, a result confirming a similar difference found in FCG mice [14]. A sex-chromosome effect on FSH levels was not found in female rats in the present study, in contrast to findings in FCG mice, in which XY females had higher FSH levels than XX females. The differential control of FSH levels as a function of sex-chromosome complement suggests that feedback control of FSH is different in XX and XY testes, despite their similar secretion of testosterone. This finding warrants further investigation to explain the mechanisms involved.

### Comparison of rat and mouse models

Although the rat models and mouse FCG model both test the differential effects of XX and XY sex chromosomes, the two models have some important differences. The mouse model utilizes a single version of ChrY, which has sustained a small deletion of the region encoding only the single-copy *Sry* gene [5]. Moreover, the *Sry* gene in the mouse model, when present, is encoded by the same transgene (which is present in multiple copies [78]), with the same copy number of *Sry* in both genotypes containing *Sry*. The XX-XY comparison is similar in gonadal males and females, except for the presence of the *Sry* transgene in the gonadal males, allowing an assessment of the differences caused by sex-chromosome complement. In the rat model, in contrast, the comparison of effects of XX and XY may be somewhat different in gonadal males and females, because of possible differences in the levels of *Sry* (Table 1). XX gonadal males may be compared to XYWT and XYTG gonadal males, all of which differ in the copy number and type of *Sry* genes. The XYWT males have the WT ChrY with potential expression of any *Sry* gene. XXTG males have expression only of the three *Sry* genes encoded by the BAC transgene. XYTG males have potential expression of *Sry* both from the WT ChrY and from the transgene. Thus, differences in phenotype of XX and XY male groups can potentially be attributed either to the XX-XY difference, or to different doses of *Sry* if the phenotype is influenced by *Sry*. Indeed, we have detected some differences in level of *Sry* expression in tissues of various genotypes (Figure 7). Similarly, when comparing in XX and XY^Δ^ gonadal female groups, a difference in phenotype may be attributable to the complement of sex chromosomes, or the expression of *Sry* from the Y^Δ^ chromosome. Some experimental outcomes reduce the probability that differences between same-gonad XX and XY groups are caused by *Sry* dose. If, for example, XYWT and XYTG groups have the same phenotype, the effect of the extra *Sry* genes in the transgene would seem not to be influencing the phenotype of interest, reducing transgene dose as the likely explanation for a difference between XXTG and XYWT rats.

The rat and mouse models differ in that the *Sry*-deficient ChrY in the mouse model encodes all non-Sry genes, whereas the CRISPR-mediated deletion of some *Sry* copies in the rat models also deleted some adjacent ChrY genes. Thus, some comparisons of groups differing only in the presence of the Y^Δ^ chromosome do not test for effects of all ChrY genes.

In the mouse model, the modified Y chromosome (“Y^−^“) is passed from the father to offspring. The loss of Sry from Y^−^ is complemented by the addition of the *Sry* autosomal transgene, so that XY^−^(*Sry*TG+) males are fully fertile. In contrast, the Y^Δ^ chromosomes produced here in rats lack some genes (probably at least *Rbmy*) required for spermatogenesis. Thus, XY^Δ^(*Sry*TG+) males are produced but have small testes and do not make sperm (data not shown). Thus, the Y^Δ^ chromosome must be passed from the mother. This necessitates two breeding schemes to produce the Four Core Genotypes of XX and XY rats with the same type of gonad. A positive ramification of passing the Y^Δ^ from the mother is that nondisjunction of X and Y^Δ^ during meiotic divisions in the ovary leads to the production of interesting sex chromosome aneuploid groups.

The mouse and rat models differ in number of genotypes produced and breeding schemes. In FCG mice, there are 4 genotypes, produced within the same litters of a single cross. Maternal or litter effects are distributed across all genotypes. The FCG cross allows measurement of sex chromosome effects (XX vs. XY) and gonad effects. The measurement of effects of number of ChrX or Chr Y are then evaluated in the XY* cross or Sex Chromosome Trisomy cross [7, 69]. In contrast, the rat models require at least two breeding schemes, one with transgenic males, and the other with an XY^Δ^ mother. Comparisons to detect effects of sex-chromosome complement (XX vs. XY, Table 1, Figure 1) are within-litter. Within each breeding cross, XX and XY rats with the same gonad are produced with only one type of gonad, and two crosses are needed to compare XX and XY rats with either type of gonad. Moreover, the comparison of rats with testes vs. ovaries (keeping sex-chromosome complement equal, either XX or XY) is made across litters. The rat crosses offer the additional advantage of producing rats with different numbers of ChrX or ChrY.

### Perspectives and Significance

Sex-chromosome effects on physiology and disease are poorly understood because specialized animal models are required to discriminate them from effects of gonadal hormones. Use of the FCG mouse models have led to the discovery of sex-chromosome effects contributing to numerous phenotypic sex differences in disease phenotypes once thought to be caused exclusively by gonadal hormones [4]. However, the limitation of these models to a single species prevents a broad understanding of a major causal mechanism of sexual differentiation. Some diseases and phenotypes are better studied in rats than mice. The rat models generated here open the way to investigation of sex-chromosome effects in a wide variety of interesting rat phenotypes in which factors protect one sex from disease more than the other sex.

## Conclusions

Our initial goal was to knock out testis-determining function on the rat ChrY, and to complement the loss of function with an autosomal testis-determining transgene. As described here, we successfully created a rat Y^Δ^ chromosome that no longer causes testis development, and generated multiple rat lines with an *Sry* transgene that successfully induces testis development in the absence of other Y genes. Together, these rat genotypes make it possible to study XX and XY rats with testes, and XX and XY rats with ovaries, which will provide unique insights into the genetic determinants of sex differences as demonstrated by numerous studies in FCG mice. The models also produce sex-chromosome aneuploid rats (XO, XXY, XYY) that are useful for measuring the dosage effects of ChrX and ChrY.

The comparison of several *Sry* transgenic lines provides the opportunity to choose a line that is most suitable for testing for sex-chromosome effects in future studies. Based on our current observations, lines 208 and 733 are most attractive. Line 424 produces smaller litters (Table 2), and involves deletion of dozens of genes at the site of integration of the transgene, considerably complicating interpretation of the effects of the transgene. Line 737 is not attractive because XXTG rats show a slightly smaller AGD relative to XYWT, suggesting that the XXTG testes do not produce as much androgen prenatally as XYWT testes. Accordingly, we no longer maintain lines 424 and 737 in our laboratories. Rat lines produced here will be available at cost from public rat repositories.

## Declarations

Ethics approvals. All experiments were approved by the Institutional Animal Care and Use Committees of participating institutions.

## Consent for publication

Not applicable

## Availability of data and materials

The datasets used during the current study are available from the corresponding author on reasonable request.

## Authors’ contributions

Conception and design of project: APA, AMG, MRD, VRH, XC, MNG, DRS, HS, DCP, KR, JWP, WG, JW. Data acquisition and analysis: APA, XC, MNG, JMR, DRS, MOT, JWP, TM, VAF, WG, LL, AT, HB, CW, LV, KR, JLC, KL, TSK, VRH, AMG; Interpretation of data: APA, XC, MNG, JMR, DRS, JWP, TM, VAF, JW, HB, CW, LV, KR, JLC, KL, TSK, VRH, MRD, AMG; drafting and revising manuscript: APA, XC, MNG, JMR, DRS, TM, VAF, WG, SL, CW, LV, HS, KR, VRH, MRD, AMG.

## Acknowledgements

Thanks to the following people for advice and assistance: Robin Lovell-Badge, James Turner, Ryohei Sekido, Haley Hrncir, Stephanie Correa, Paul E. Micevych, Laura Cortes, Margaret Mohr, Nancy Forger, Stuart Tobet, Margaret M. McCarthy, Jan Willing, Mansoureh Eghbali, Kathryn Sandberg, Julia Gong, Morgyn Michelson, Edith Heard, James Cleland, Agnese Loda, Yuichiro Itoh, Thom Saunders and staff of the University of Michigan Animal Model Core, University of Virginia Ligand Assay & Analysis Core, and the Monash University Histology Platform.

---

**Supplementary Figure 1.**
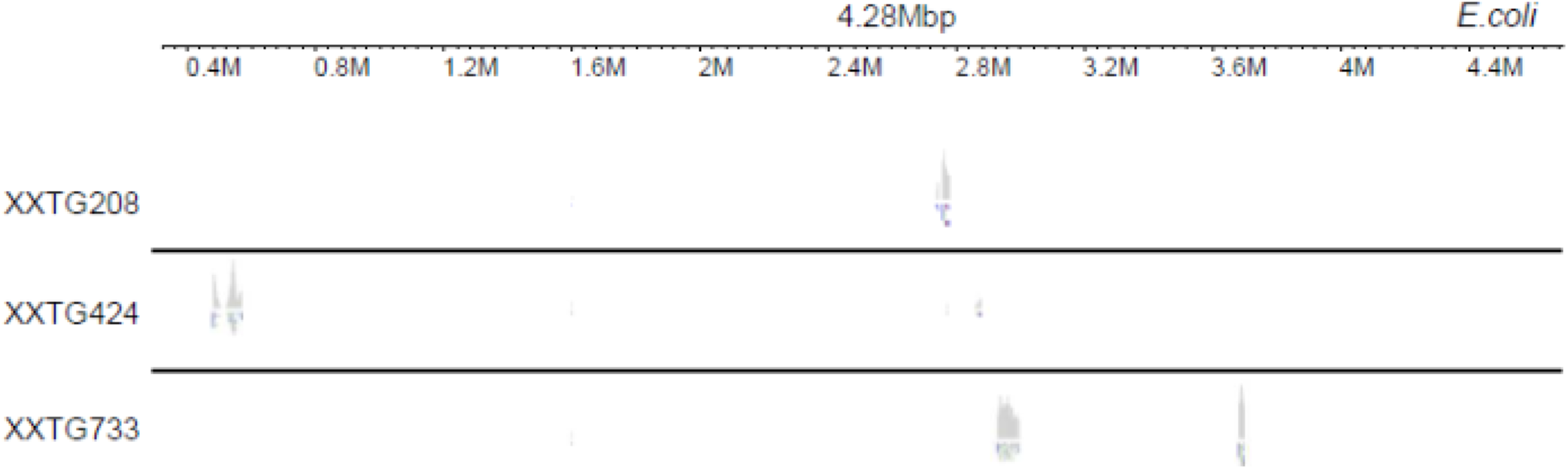
Alignment of long-read sequence to laboratory host *E. coli* bacterial genomic DNA. Some *E. coli* genomic DNA copurified with the BAC prior to microinjection is co-inserted at each integration site within each transgenic line. Aligned regions ranged from ∼2.4- to ∼68-kbp.

**Supplementary Figure 2:**
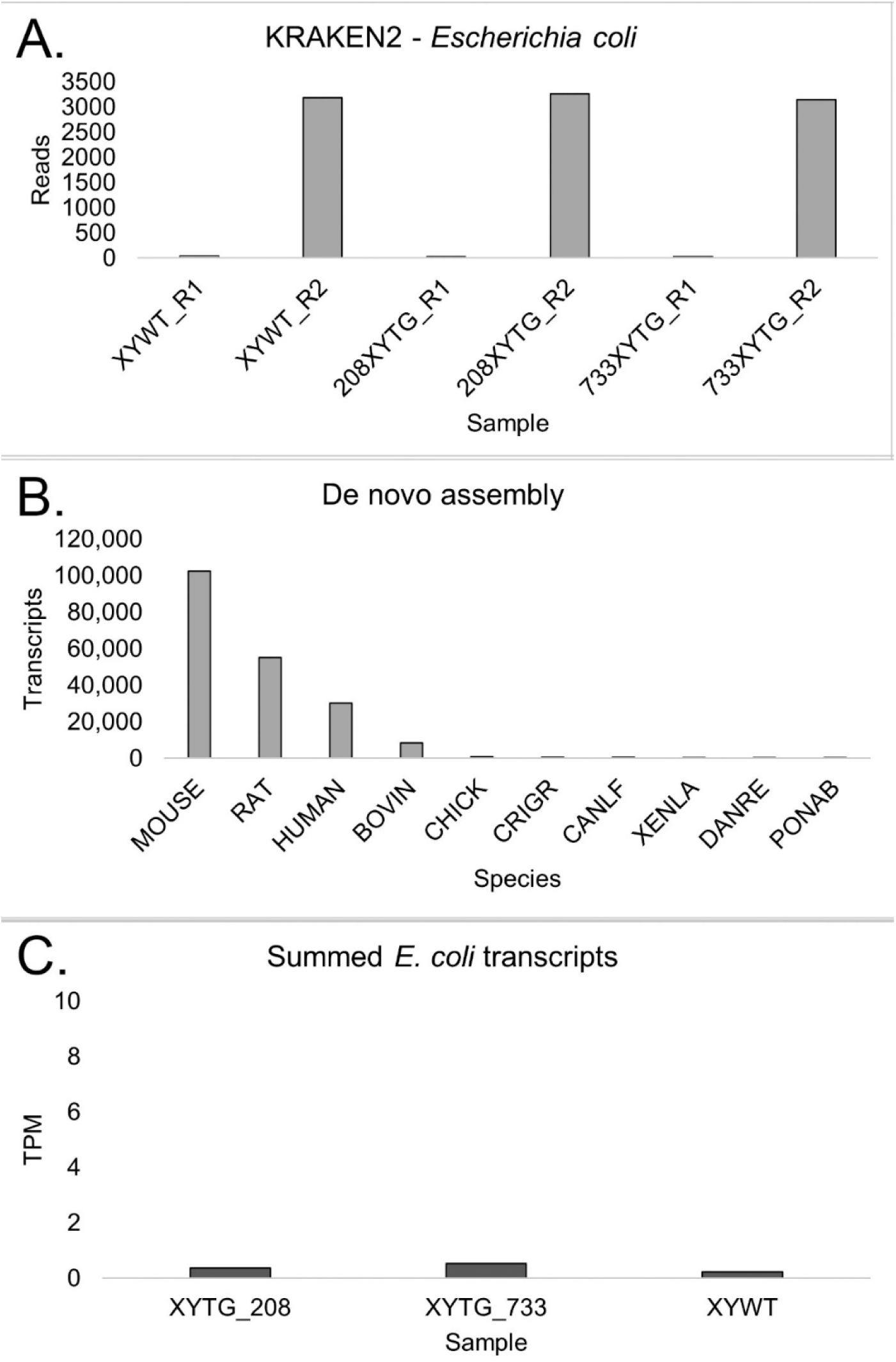
Testing for expression of *E. coli* transcripts A. The number of reads mapping to *E coli* genome from either end (R1 or R2) of liver RNAseq per sample. The read numbers were similar in two transgenic lines 208 and 733, which contain some *E. coli* genome, relative to an XYWT male, which lacks any E. coli genome, indicating that read numbers were at background in each case. B. Number of transcripts per species from de novo assembly. BOVIN, bovine; CRIGR, *Cricetulus griseus* (Chinese hamster); CANLF, *Canis lupus familiaris* (dog); XENLA, *Xenopus laevis*; DANRE, *Danio rerio;* PONAB, *Pongo abelii* (Sumatran orangutan). C. Transcripts per million (TPM) of the three *de novo* constructed *E. coli* transcripts were all below 1 TPM background noise.

